# Mosaic Regulation of Stress Pathways Underlies Senescent Cell Heterogeneity

**DOI:** 10.1101/2024.10.03.616489

**Authors:** Roberto A. Avelar, Thomas Duffield, Cyril Lagger, Nikita Krstevska, Marian Breuer, João Pedro de Magalhães

## Abstract

Cellular senescence (CS) and quiescence (CQ) are stress responses characterised by persistent and reversible cell cycle arrest, respectively. These phenotypes are heterogeneous, dependent on the cell type arrested and the insult inciting arrest. Because a universal biomarker for CS has yet to be identified, combinations of senescence-associated biomarkers linked to various biological stress responses including lysosomal activity (β-galactosidase staining), inflammation (senescence-associated secretory phenotypes, SASPs), and apoptosis (senescent cell anti-apoptotic pathways) are used to identify senescent cells.

Using in vitro human bulk RNA-seq datasets, we find that senescent states enrich for various stress responses in a cell-type, temporal, and insult-dependent manner. We further demonstrate that various gene signatures used to identify senescent cells in the literature also enrich for stress responses, and are inadequate for universally and exclusively identifying senescent samples.

Genes regulating stress responses – including transcription factors and genes controlling chromatin accessibility – are contextually differentially expressed, along with key enzymes involved in metabolism across arrest phenotypes. Additionally, significant numbers of SASP proteins can be predicted from senescent cell transcriptomes and also heterogeneously enrich for various stress responses in a context-dependent manner.

We propose that ‘senescence’ cannot be meaningfully defined due to the lack of underlying preserved biology across senescent states, and CS is instead a mosaic of stress-induced phenotypes regulated by various factors, including metabolism, TFs, and chromatin accessibility. We introduce the concept of Stress Response Modules, clusters of genes modulating stress responses, and present a new model of CS and CQ induction conceptualised as the differential activation of these clusters.

## 1. Introduction

Cellular senescence (CS) – often characterised as irreversible cell cycle arrest – influences ageing, tumour suppression, tumorigenesis, chronic diseases, wound healing, regeneration, embryonic receptivity, and development [1–9]. CS is induced via replicative senescence (RS) due to telomere erosion [10–12], stress-induced premature senescence (SIPS) by DNA and cellular damage [13–17], and oncogene-induced senescence (OIS) through erroneous oncogene activation [18–20], among other stressors.

There are various issues with how CS is defined, and identifying senescent cells is challenging due to the lack of universal biomarkers; a multifaceted approach using various biomarkers is required, complicating senescence detection, particularly in vivo [21–23]. Biomarkers include cell cycle arrest, β-galactosidase (β-gal) staining, senescence-associated secretory phenotype (SASP) secretions, senescence-associated heterochromatin foci (SAHF) formation, and an enlarged and flattened cellular morphology, plus the expression of cyclin-dependent kinase inhibitors and tumour-suppressor genes like p53, p21, and p16 [22]. However, all of these biomarkers can be uncoupled from CS and are associated with other processes (Table 1).

**Table 1.** Biomarkers of CS alongside examples from the literature highlighting how these biomarkers have been uncoupled from senescence and are detectable in contexts outside of CS, indicating that there is no universal, process-specific biomarker of CS.

| Senescence Biomarker | Uncoupled from Senescence | Relevant to other Processes |
| --- | --- | --- |
| β-galactosidase staining | Knockdown of *GLB1* leads to senescence-associated cell cycle arrest without β-gal staining [24]. | Activated macrophages and quiescent cells both stain positive for β-gal, a biomarker of lysosomal stress [25]. |
| Secretory phenotypes | Knockdown of *BRD4* blunts SASP gene expression in OIS and SIPS even after cells have established a senescence phenotype [26]. Additionally, mouse cells induced into CS at 20% oxygen concentration lack a SASP despite being irreversibly arrested [27, 28]. | Activated and cancer-associated fibroblasts secrete various SASP-associated factors, including VEGFs, cytokines like IL6 and IL8, chemokines, and matrix metalloproteinases, while maintaining the ability to proliferate [29]. Immune and endothelial cells also secrete factors reminiscent of the SASP [22]. |
| Hypophosphorylation of Rb protein | Activation of the p53-dependent DNA damage response can trigger senescence in cells with dysfunctional Rb protein [30]. | Rb protein is hypophosphorylated in CQ [31]. |
| Senescence-associated heterochromatin foci | SAHF are dispensable for cellular senescence and primarily associated with OIS and not RS or SIPS [32]. | DNA damage repair-deficient oncogene-expressing cells have nuclear heterochromatic structures morphologically reminiscent of SAHF while maintaining their proliferative capacity [33]. |
| p53/p21 and p16 expression | Cells can be induced into SAHF-dependent irreversible arrest independent of p16, p53, and p21 via downregulation of p300 histone acetyltransferase [34]. | p53 and p21 expression are implicated in other cell cycle arrest phenotypes besides CS, including CQ and terminal differentiation [35-38]. |
| Irreversible arrest | Senescent BJ fibroblasts can re-enter the cell cycle following p53 knockdown, maintaining SASP secretions [39]. | Granulocyte-monocyte ER-HOXA9 cells lose their ability to proliferate following terminal differentiation into mature neutrophils and monocytes [40]. |
| DNA damage | Senescence phenotypes can be triggered without detectable DNA damage [41]. | Quiescent hematopoietic stem cells accumulate DNA damage that is repaired upon re-entry into the cell cycle [42]. |
| Enlarged, flattened morphology | Enforcing senescent cells to have a spindle-shaped morphology – as opposed to | TGF-β-treated prostate epithelial cells exhibit elevated SA-β-gal activity alongside CS morphology whilst |
|  | the typical enlarged, flattened morphology – does not allow re-entry into the cell cycle [43]. Furthermore, RAF-induced OIS results in ‘retracted spindle’ or ‘spherical’ morphologies as opposed to traditional CS-associated morphologies [44]. | simultaneously maintaining their proliferative capacity [45]. |

Senescent cells exhibit heterogeneity in both phenotype and function [46, 47]. The composition of the SASP is heterogeneous, dependent on insult and genetic factors such as p53, RAS, and p16 [27, 48]. The transcriptomic profile of senescent cells and their respective SASPs are partially dependent on the cell type, insult, and physiological environment of the cell when CS is induced [27, 47, 49]. This heterogeneity suggests that senescent cells tailor their functions to their biological context, as seen in senescent pancreatic β-islets which secrete more insulin than their non-senescent counterparts [50]. Furthermore, post-mitotic cells like neurons and muscle cells showcase senescence-associated features under stress – like SASP production – despite lacking proliferative potential to begin with [51, 52].

The reversibility of arrest phenotypes in CS [39, 44, 53, 54] is itself contradictory; CS is classically defined as irreversible cell cycle arrest. SAHF – which are p16-dependent and primarily associated with OIS [32] – contribute to CS-associated irreversible arrest phenotypes [34, 55, 56]. On the other hand, p53/p21, known to induce CS [57], are also implicated in post-mitotic terminal differentiation and CQ – cellular programmes that showcase more-readily reversible forms of arrest [36, 37, 58]. The role of p53 and p21 in maintaining these states – alongside the fact that arrest associated with p53-induced CS can be reversed – blurs the line between ‘irreversible’ arrest in CS compared to reversible arrest in other contexts. Moreover, CQ is not a single uniform state, and ‘deeper’ quiescent depths are implicated in the transition from CQ into CS [31, 59, 60]. As such, at least two mechanisms of cell cycle arrest – a more readily reversible arrest associated with p53/p21 expression compared to a stringent irreversible arrest associated with p16 expression and erroneous oncogene activation – appear to have evolved in mammals.

CS manifests as a gradual process; a sequential emergence of gain-of-senescence phenotypes associated with specific genetic clusters has been identified in RS [61–63]. Secretion of SASP factors is also accelerated in OIS compared to other CS phenotypes [27, 64]. Uncoupling of the SASP from cell cycle arrest further indicates distinct regulatory mechanisms between these processes. Indeed, the regulation of SASP secretions varies, with a greater involvement of chromosomal rearrangements in OIS compared to RS, possibly via mechanisms involving SAHF [65, 66]. Furthermore, metabolic alterations such as in the prostaglandin pathway have been shown to drive SASP heterogeneity [49, 67–70]. The temporal nature of CS – alongside the uncoupling of CS biomarkers from senescent states – suggests that senescence itself is a combination of multiple phenotypes [62, 71, 72]. Furthermore, p53 is involved in various other stress responses, including CQ, DDR signalling, autophagy, inflammation, and apoptosis, indicating that CS phenotypes may encompass multiple stress responses [35, 57, 64, 73–82].

In this study, we perform a bioinformatic analysis of CS and CQ transcriptomes and find that transcriptomic markers of CS commonly used to identify senescent cells in the literature fail to do so in a universal and exclusive manner. Furthermore, CS and CQ transcriptomes encompass various stress response pathways, including lysosomal genes, inflammation, apoptosis, and hypoxia. We further show transcriptomic heterogeneity of TFs, metabolic enzymes, epigenetic regulators, and key stress response genes that potentially heterogeneously regulate stress response pathways in CS. As such, we suggest that heterogeneity observed in mammalian cell cycle arrest phenotypes is due to differential regulation of stress responses, which do not universally coactivate alongside reversible and irreversible proliferative arrest. We call the clusters of genes associated with separate CS biomarkers ‘Stress Response Modules’ (SRMs) (Figure 1). This model suggests that senescent cell heterogeneity is due to mosaic activation of tailored stress-associated pathways, with CS not distinctly classifiable as a specific subset of SRMs or any other discrete category.

**Figure 1.**
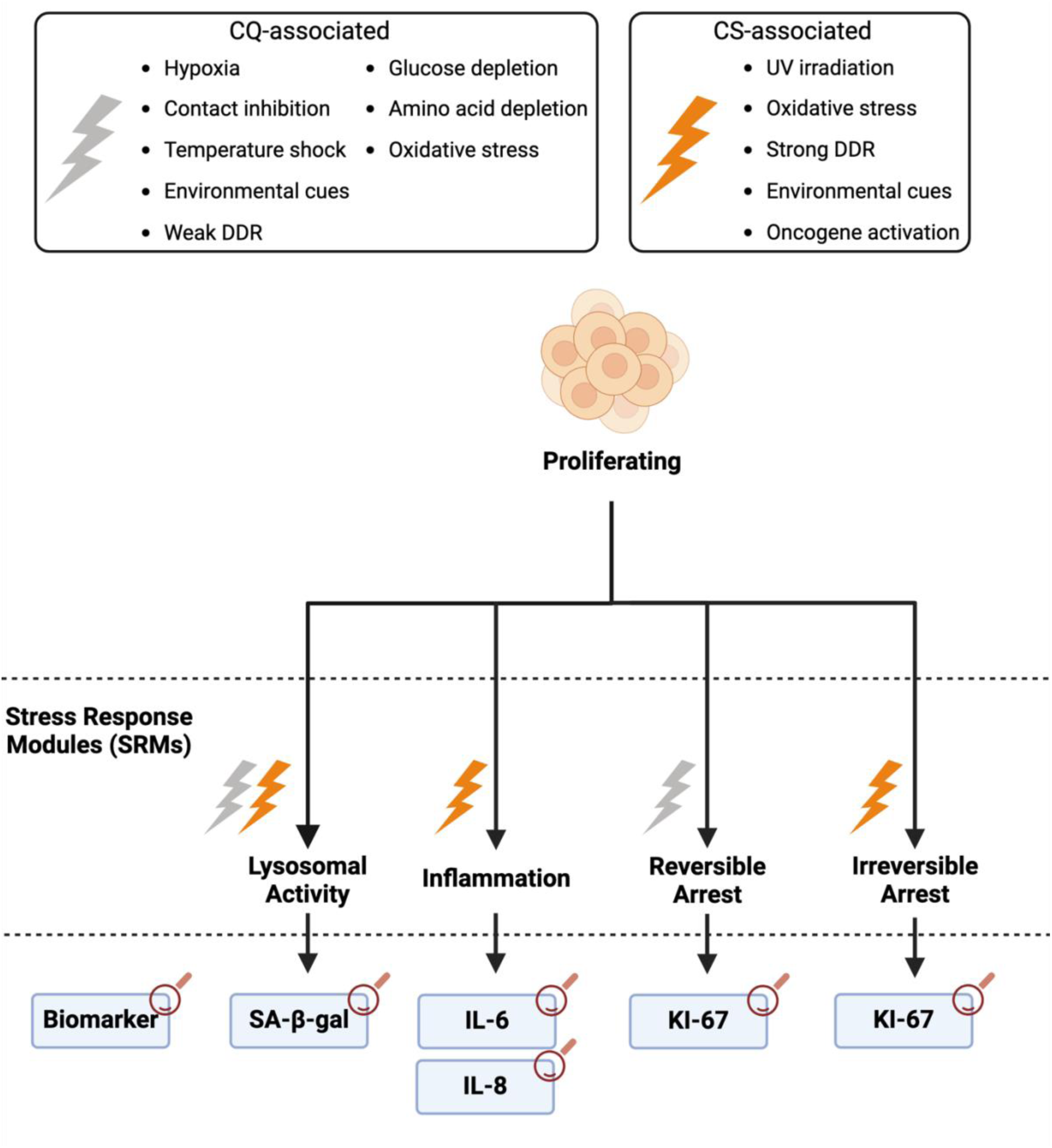
Proposed model of senescence and quiescence induction as differential activation of stress response modules. The activation of different modules is currently measured via different biomarkers, including β-gal as a proxy marker of lysosomal activation associated with autophagy, while inflammatory genes are used as markers for the activation of secretory phenotypes [22]. Importantly, activation of SRMs can occur independent of cell cycle arrest; the coactivation of these modules is not guaranteed in senescence, and individual stress response phenotypes like inflammation and lysosomal activity have been uncoupled from cell cycle arrest in ‘senescent’ cells [24, 27, 83, 84]. Moreover, distinct classes of cell cycle arrest modulated by the p53/p21 and p16 pathways likely influence the reversibility of SRM activations [39].

## 2. Results

### 2.1 Transcriptional Heterogeneity of Arrested Human Fibroblast Cell Lines

We probed the senescent, quiescent, and proliferating transcriptomic profiles of human lung, skin, and foreskin fibroblast samples across 34 studies, focusing on uniformly processed bulk RNA-seq datasets from recount3 [85, 86] (SI Table 1). OIS samples included cells expressing H-RasV12 (HRAS) (n=42) or BRAFV600E (BRAF) (n=6) constructs, or corresponding control samples transfected with control siRNAs (see 5.1 Cell Cycle Arrest Transcriptomic Data). SIPS samples were induced into CS via DNA damage, while CQ was induced via contact-inhibition (n=19) or serum-starvation (n=22), and RS was induced via proliferative exhaustion.

#### 2.1.1 Cell Cycle Arrest Transcriptome Comparison

After removing the study batch effect via linear regression, principal component analysis (PCA) was performed to assess how samples clustered (SI Figure 1); the top two PCs accounted for 71% of sample variation. We identified 4 separate clusters: i) proliferating; ii) serum-starved and contact-inhibited CQ; iii) SIPS and RS; and iv) OIS. Differentially expressed genes (DEGs) were derived between arrested samples and proliferating controls using DESeq2 (p<0.05 and |log_2_FC|>log_2_(1.5), negative binomial distribution with Benjamini-Hochberg (BH) false discovery rate (FDR) correction) [87] (SI Table 2, see 5.2 in methods). Volcano plots of DEGs are shown in SI Figure 2. When we considered genes showcasing the largest variance across samples, the top genes consisted of SASP factors including *IL1B*, *MMP3*, *CXCL8*, and *SERPINB2* (SI Table 2).

Across all five cell cycle arrest states, there were 316 and 101 shared under- and overexpressed DEGs respectively (SI Figure 3, SI Table 3). We performed 10,000 simulations to determine the likelihood of DEGs changing in the same direction across all conditions (SI Table 4). Across simulations, DEGs never changed in the same direction by chance more than 13 times (see 5.4 in methods) (SI Figure 4, SI Table 5) indicating that there are significantly more arrest-DEGs shared between arrested conditions than expected.

The shared under- and overexpressed DEGs were enriched using genes that change in the same direction in all five arrest conditions – regardless of significance – as an enrichment background (SI Table 6-7). There were no enriched KEGG or GO terms for shared overexpressed DEGs. Unsurprisingly, the shared underexpressed DEGs enriched for cell cycle-associated terms, including ‘cell cycle,’ ‘cell division,’ and ‘meiotic cell cycle process’ (SI Figure 5a). DNA repair pathways were also enriched, alongside pathways involved in response to irradiation. The ‘Cellular senescence’ KEGG pathway was enriched amongst the shared underexpressed DEGs (SI Figure 5b), although the KEGG CS pathway constitutes both genes that promote and inhibit proliferation; amongst shared underexpressed DEGs are cyclin A2, B1, B2, and CDK1, which are necessary for cell cycle progression and are expected to downregulate in arrest.

Samples were grouped via unsupervised hierarchical clustering based on the top 15 over- and underexpressed DEGs (identified using π scores) for each cell cycle arrest condition (SI Figure 6, SI Table 8) (see 5.3 PCA and Heatmaps). All proliferating, CQ, and CS samples clustered into their respective groups. Nonetheless, these DEGs were unable to fully differentiate between CQ subgroups.

Over-representation analysis (ORA) was performed between conditions using all genes expressed within the recount3 data as an overlap background, and significantly underexpressed DEGs were significantly shared across arrest conditions (p<0.05, two-tailed Fisher’s exact test with Bonferroni correction) (Figure 2a, SI Figure 7, SI Table 9). This was also the case with the overexpressed arrest DEGs. The exception was the overlap between overexpressed OIS DEGs with overexpressed contact-inhibited CQ DEGs. The overexpressed CQ DEGs did not significantly overlap with the underexpressed CQ DEGs, as expected. We found the same pattern with the SIPS DEGs, where the overexpressed CQ DEGs overlapped the underexpressed SIPS DEGs significantly less than expected by chance while the underexpressed CQ DEGs overlapped the overexpressed SIPS DEGs less than expected by chance. However, it appears that more overexpressed RS and OIS DEGs are underexpressed in serum-starved CQ than expected by chance.

**Figure 2.**
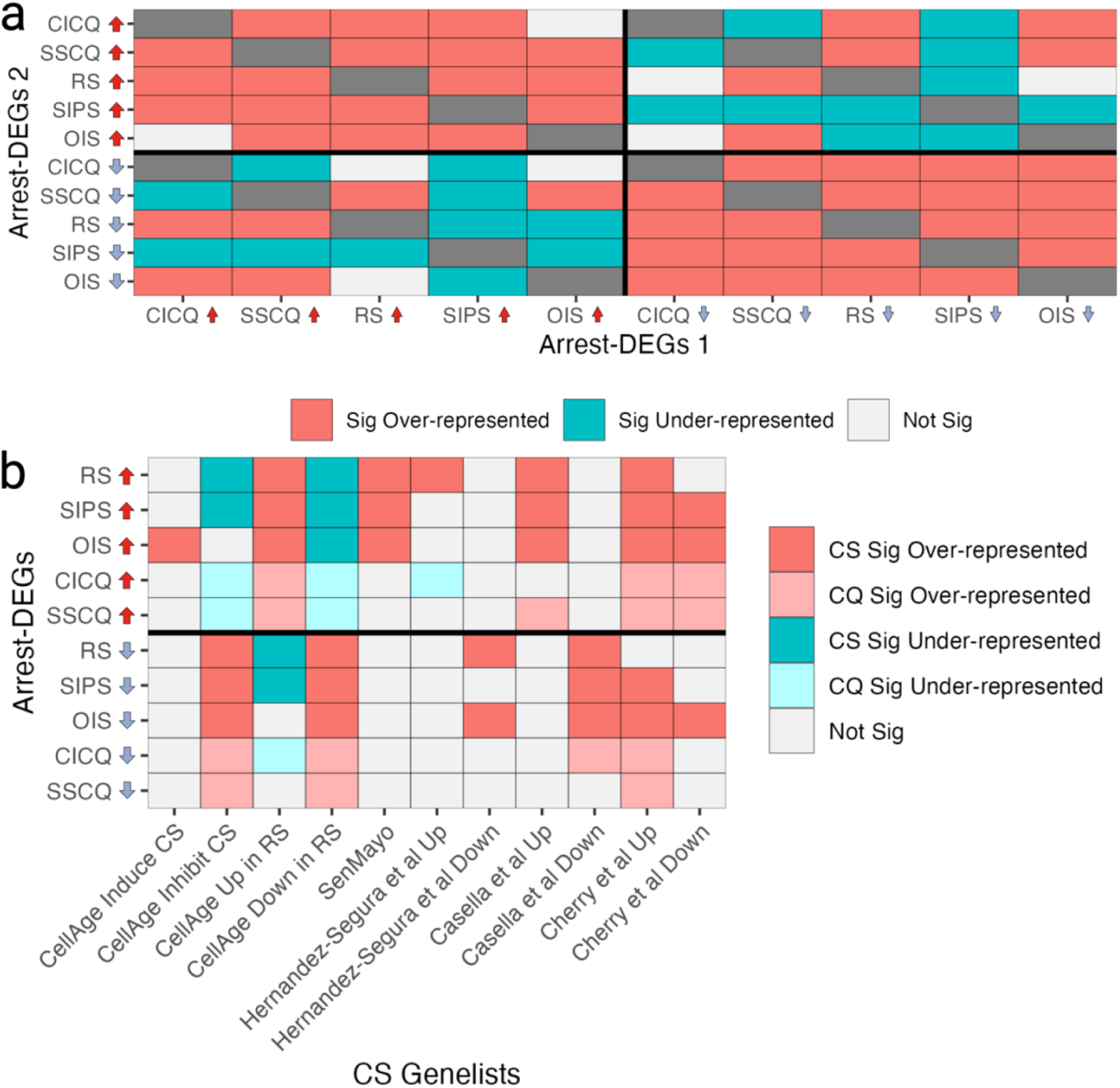
a) Overlap between arrest-DEGs. b) Overlap of arrest-DEGs with various gene lists of interest. Red tiles indicate significantly more overlaps than expected by chance, whereas blue tiles indicate significantly fewer overlaps than expected, and only significant results are shown (p<0.05, two-tailed Fisher’s exact test with Bonferroni correction). For (b) lighter tiles are used to distinguish quiescence from senescence.

Various transcriptomic signatures of CS have been published (Table 2). To determine whether these signatures are significantly associated universally with CS – and not CQ – we overlapped them with the arrest DEGs using genes expressed in the recount3 fibroblast data as the background (Figure 2b, SI Table 10-11). We also overlapped CellAge genes that are capable of inducing and inhibiting CS when genetically perturbed.

**Table 2.** Sources of CS signatures from various published studies. Genes available in SI Table 10.

| Signatures | Explanation |
| --- | --- |
| CellAge Drivers | Genes that induce or inhibit the senescence phenotype when genetically manipulated in human cell lines [21, 88]. |
| CellAge signatures of RS | Signatures of RS compiled from a meta-analysis of human cells [89]. |
| SenMayo | Gene set used to identify senescent cells across tissues and species [90]. |
| Hernandez-Segura et al | Senescence-associated 'core' signatures identified in senescent human fibroblasts, melanocytes, astrocytes, and keratinocytes and validated in mouse cells, generated via irradiation and oxidative stress [47]. From the data generated in house, only samples sequenced 10 days post-irradiation were included in the generation of the 'core' signatures of CS. |
| Casella et al | Senescence signatures developed from human fibroblasts and endothelial cells induced into CS via RS, ionising radiation, doxorubicin, or HRASG12V overexpression [91]. |
| Cherry et al | In vivo derived signature of CS from p16<sup>+</sup> fibrotic mice validated in human scRNA-seq datasets [92]. The authors themselves note that these signatures are not universal across tissues. |

We assessed whether any of these signatures could be used as universal transcriptomic signatures of CS, based on the following criteria: i) be significantly overrepresented in all senescent conditions – in a direction-dependent manner where applicable – and ii) be unique to CS as opposed to other biological processes like CQ. The majority of these gene lists failed to meet these criteria.

The most promising gene list that was significantly upregulated exclusively in CS and not CQ was SenMayo. However, when we looked at the genes that were shared exclusively across CS conditions, we only found 10 genes upregulated across CS – *ANGPTL4*, *CCL26*, *CSF2*, *CST4*, *EREG*, *FGF2*, *MMP12*, *MMP3*, *NRG1*, and *SPX* – indicating that the majority of overexpressed SenMayo genes are not universally shared exclusively in CS (SI Figure 8, SI Table 12).

#### 2.1.2 Associations Between Cell Cycle Arrest and Stress Responses

To find potential SRMs in arrest phenotypes, ORA was performed between arrest-DEGs and all gene lists from the Molecular Signatures Database (MSigDB) hallmark gene set collection. Furthermore, given associations between arrest phenotypes and lysosomal activity, we included a list of lysosome-related genes published by Bordi et al. [93] (SI Table 13).

There was significant underexpression of mitotic spindle, G2M checkpoint, and E2F target genes across arrest conditions, as expected (Figure 3a, SI Figure 9a, SI Table 14). Furthermore, lysosomal genes in arrest phenotypes were significantly overexpressed, except in serum-starved CQ. While overlaps suggest that all CS conditions are significantly associated with inflammation, there was heterogeneity amongst which proinflammatory pathways were upregulated in CS, with the interferon gamma response and IL6 JAK STAT3 signalling pathways overexpressing in OIS but not SIPS or RS. Finally, both the apoptosis and p53 pathways were significantly overrepresented amongst overexpressed CS – but not CQ – DEGs.

**Figure 3.**
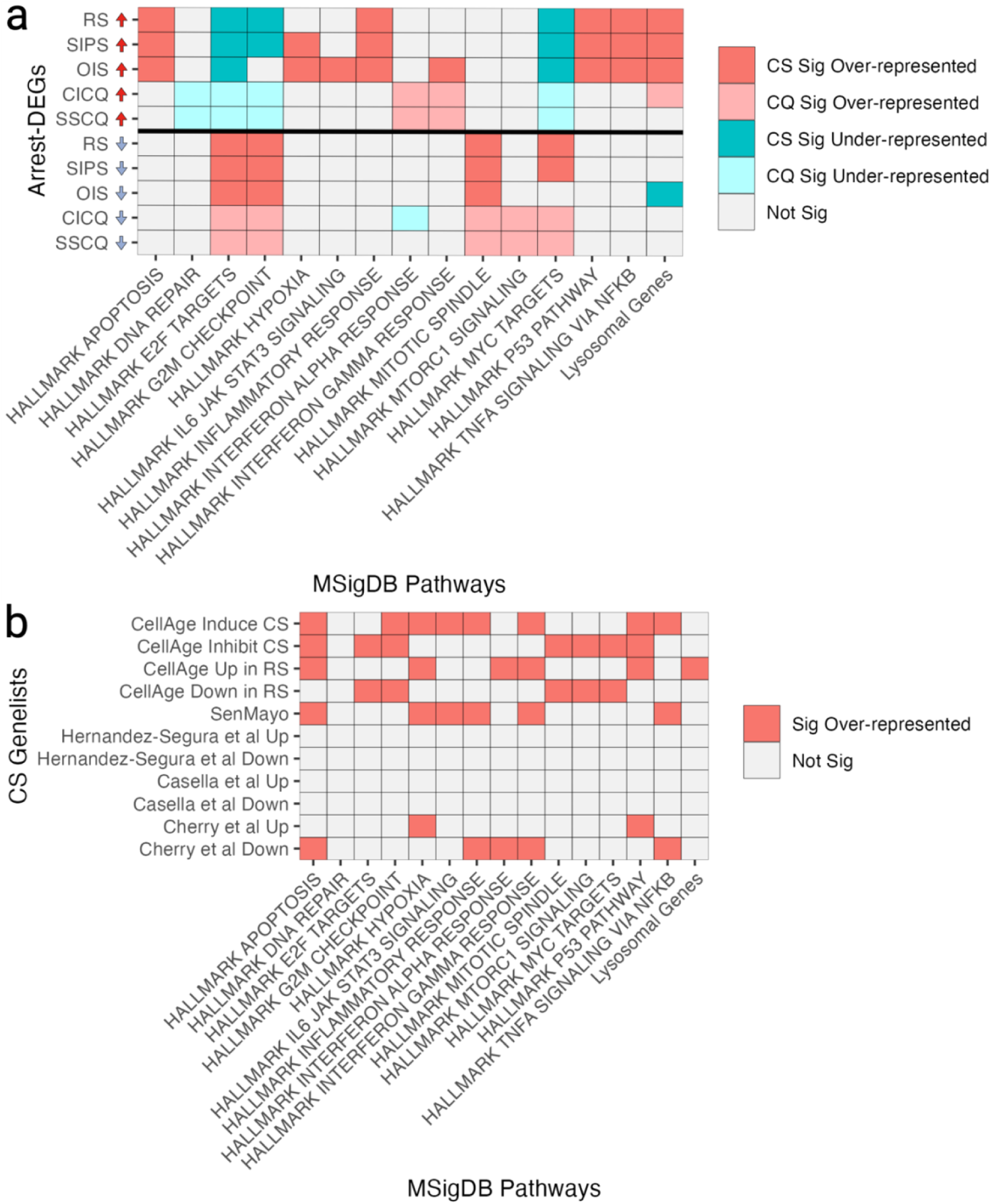
a) Overlap between arrest-DEGs and various pathways of interest. Red tiles indicate significantly more overlaps than expected by chance, whereas blue tiles indicate significantly fewer overlaps than expected. b) Overlap between various gene lists and pathways of interest. Note that while overlaps were performed for the entire MSigDB, only some pathways of interest are shown due to space constraints.

We considered whether senescence gene lists might also capture stress responses. ORA was performed between the gene lists from Table 2 and MSigDB, using genes expressed in the fibroblast data as a background for consistency (Figure 3b, SI Figure 9b, SI Table 15). Overexpressed CellAge RS signatures significantly overlapped the lysosomal genes, alongside the p53 pathway, various inflammation pathways, hypoxia, and apoptosis pathways. Underexpressed CellAge RS signatures were associated with MYC targets, MTORC1 signalling, and proliferation-associated pathways. CellAge genes were also associated with various stress responses and pathways. Finally, SenMayo and the Cherry et al. datasets are also measuring various stress response pathways including hypoxia, inflammation, and apoptosis, although not all gene lists enrich for stress responses.

Given the connections between SenMayo and stress pathways, we questioned whether other cellular states might also enrich for SenMayo. Literature indicates that senescent cells exhibit behaviour similar to activated macrophages, characterised by increased secretory and lysosomal activity [94]. To explore this, we compared SenMayo with DEGs generated from both classically and alternatively activated macrophages against untreated macrophages [95] (SI Figure 10, SI Table 10 and 16). Since the macrophage data is from mice, we conducted ORA using the SenMayo mouse gene list— approximately 80% of which has an equivalent human homologue from the human SenMayo list — using all protein-coding mouse genes as the background. SenMayo was significantly overrepresented amongst both classes of activated macrophages, indicating that while SenMayo is significantly enriched exclusively in the CS DEGs, it does not separate CS from other stress-related phenotypes.

To assess whether there is crosstalk between the MSigDB pathways, ORA was performed between different pathways using genes expressed in the fibroblast data as a background for consistency (SI Figure 11, SI Table 17). Various genes are shared across pathways, like the apoptosis gene list which significantly overlaps the hypoxia pathway, various pro-inflammatory pathways, and the MTORC1 pathway. As such, we assessed the expression of individual genes linked to various stress pathways and processes related to CS and CQ based on the literature (Table 4). We focused on genes known to promote or inhibit apoptosis, alongside genes associated with autophagy and lysosome function, inflammation, cell cycle arrest, chromatin architecture in SAHF formation, and transcription factors (TFs) associated with regulating stress responses, and found widespread heterogeneity dependent on insult (Figure 4a).

**Figure 4.**
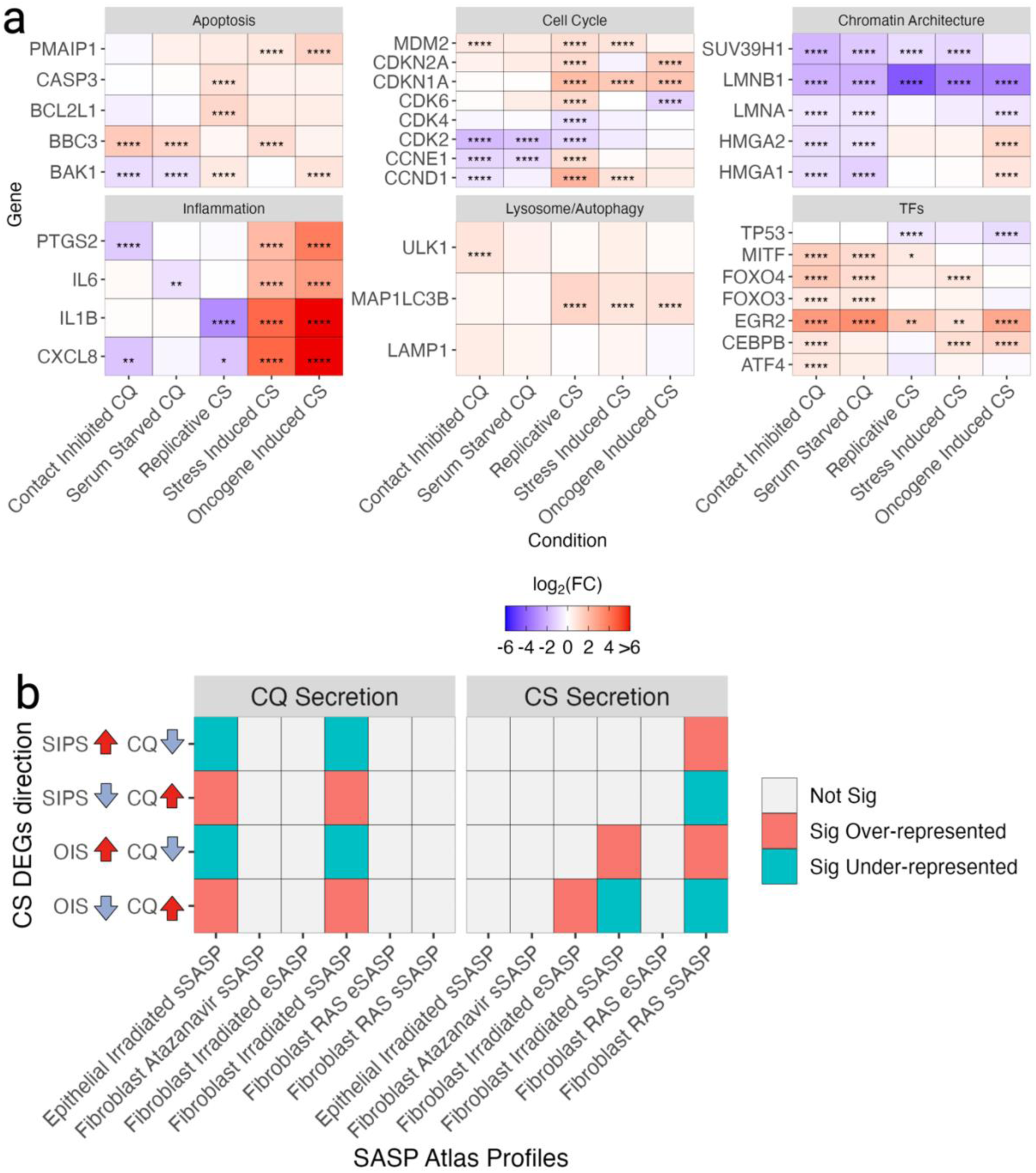
a) Heterogeneous expression of key stress response genes across cell cycle arrest conditions, compared to proliferating controls. Red tiles indicate overexpression of the given genes for the cell cycle arrest conditions compared to proliferating controls, while blue tiles indicate underexpression. Significance assessed using a negative binomial distribution with BH correction and |log_2_(FC)|>log_2_(1.5)). Maximum log_2_FC was capped at 6 to visualise differences more clearly between conditions. b) Overlap of DEGs generated between CS and CQ samples and SASP secretomes.

**Table 4.**
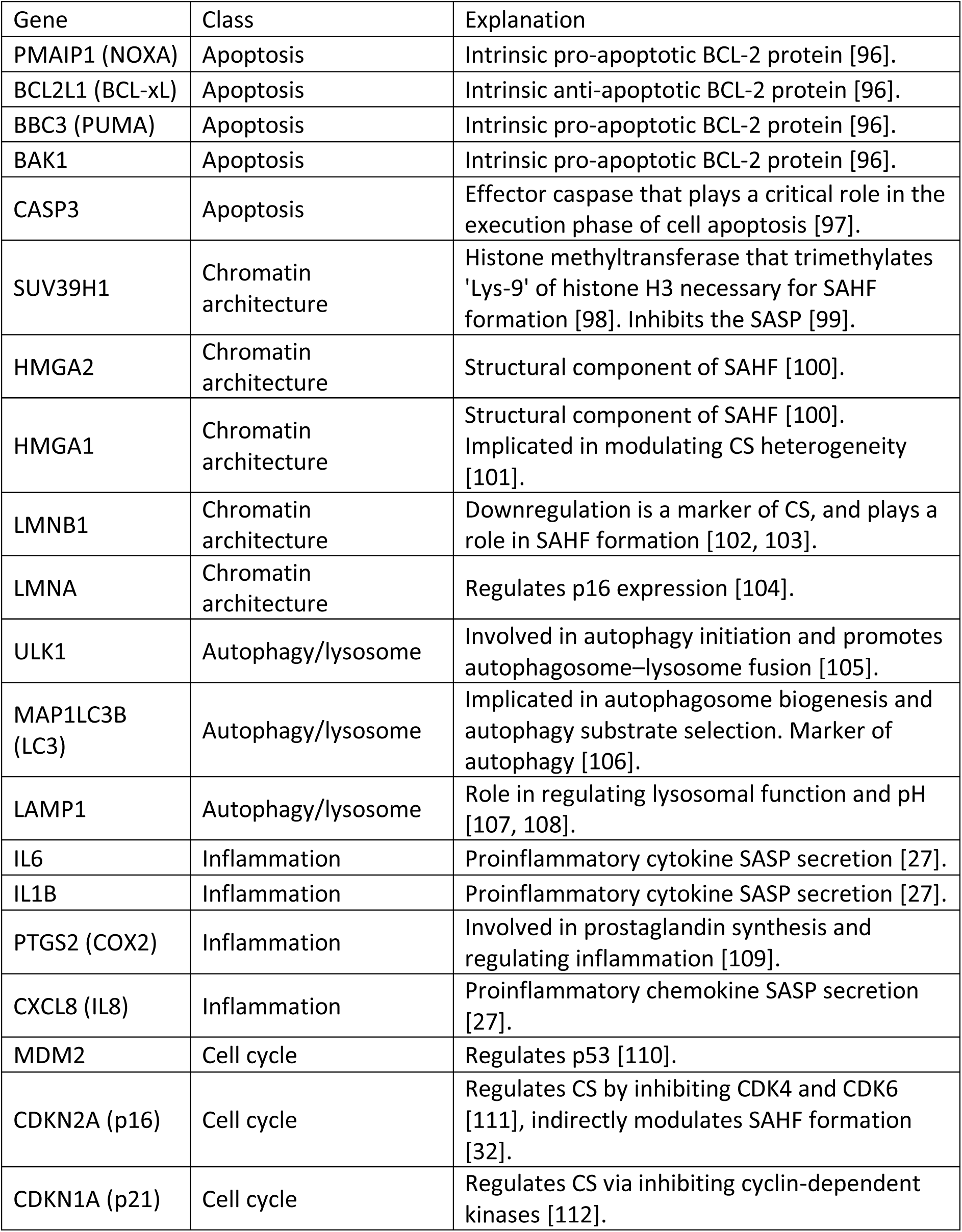
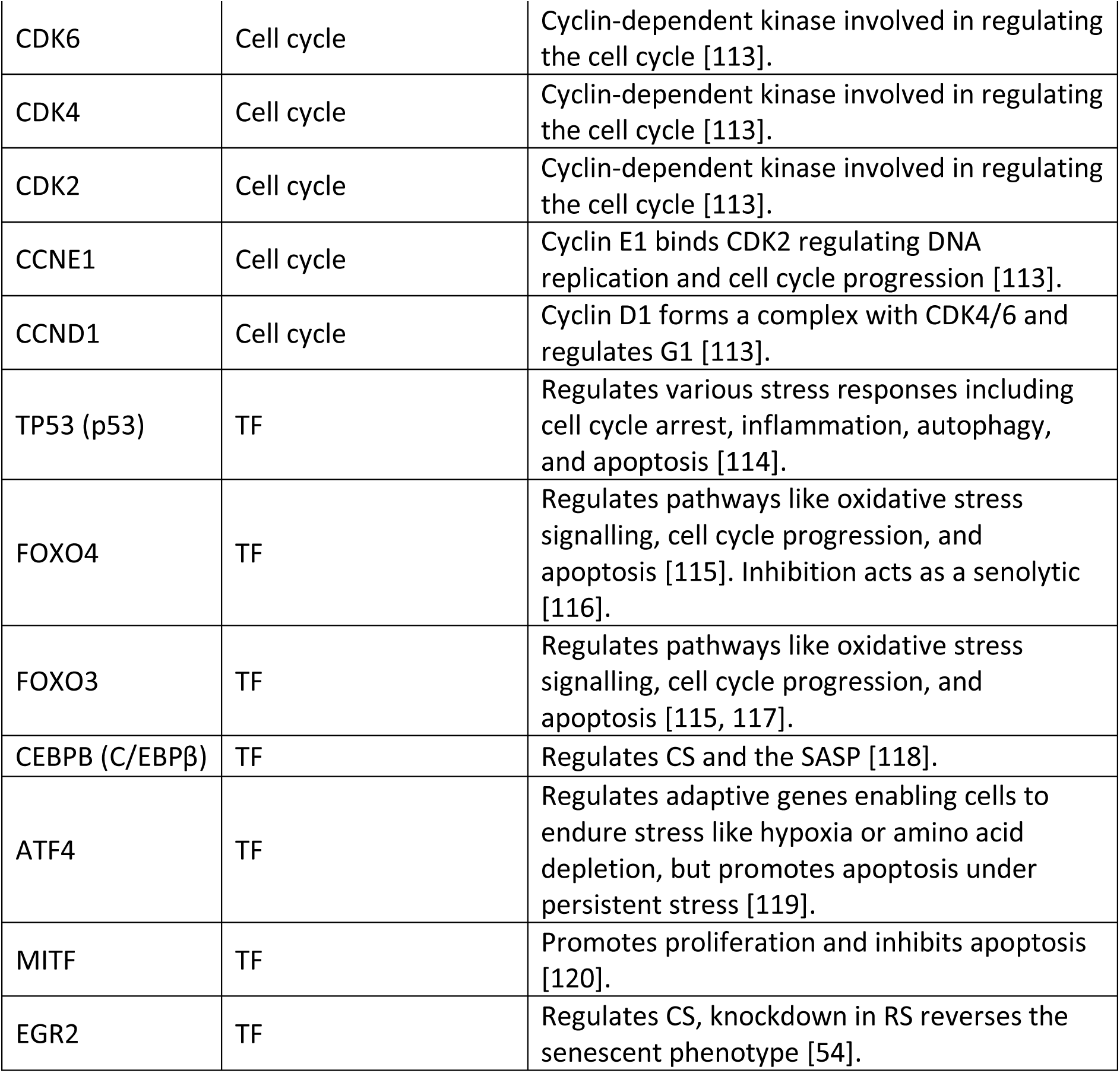
Various genes associated with CS or other stress response pathways.

| Gene | Class | Explanation |
| --- | --- | --- |
| PMAIP1 (NOXA) | Apoptosis | Intrinsic pro-apoptotic BCL-2 protein [96]. |
| BCL2L1 (BCL-xL) | Apoptosis | Intrinsic anti-apoptotic BCL-2 protein [96]. |
| BBC3 (PUMA) | Apoptosis | Intrinsic pro-apoptotic BCL-2 protein [96]. |
| BAK1 | Apoptosis | Intrinsic pro-apoptotic BCL-2 protein [96]. |
| CASP3 | Apoptosis | Effector caspase that plays a critical role in the execution phase of cell apoptosis [97]. |
| SUV39H1 | Chromatin architecture | Histone methyltransferase that trimethylates 'Lys-9' of histone H3 necessary for SAHF formation [98]. Inhibits the SASP [99]. |
| HMGA2 | Chromatin architecture | Structural component of SAHF [100]. |
| HMGA1 | Chromatin architecture | Structural component of SAHF [100]. Implicated in modulating CS heterogeneity [101]. |
| LMNB1 | Chromatin architecture | Downregulation is a marker of CS, and plays a role in SAHF formation [102, 103]. |
| LMNA | Chromatin architecture | Regulates p16 expression [104]. |
| ULK1 | Autophagy/lysosome | Involved in autophagy initiation and promotes autophagosome–lysosome fusion [105]. |
| MAP1LC3B (LC3) | Autophagy/lysosome | Implicated in autophagosome biogenesis and autophagy substrate selection. Marker of autophagy [106]. |
| LAMP1 | Autophagy/lysosome | Role in regulating lysosomal function and pH [107, 108]. |
| IL6 | Inflammation | Proinflammatory cytokine SASP secretion [27]. |
| IL1B | Inflammation | Proinflammatory cytokine SASP secretion [27]. |
| PTGS2 (COX2) | Inflammation | Involved in prostaglandin synthesis and regulating inflammation [109]. |
| CXCL8 (IL8) | Inflammation | Proinflammatory chemokine SASP secretion [27]. |
| MDM2 | Cell cycle | Regulates p53 [110]. |
| CDKN2A (p16) | Cell cycle | Regulates CS by inhibiting CDK4 and CDK6 [111], indirectly modulates SAHF formation [32]. |
| CDKN1A (p21) | Cell cycle | Regulates CS via inhibiting cyclin-dependent kinases [112]. |
| CDK6 | Cell cycle | Cyclin-dependent kinase involved in regulating the cell cycle [113]. |
| CDK4 | Cell cycle | Cyclin-dependent kinase involved in regulating the cell cycle [113]. |
| CDK2 | Cell cycle | Cyclin-dependent kinase involved in regulating the cell cycle [113]. |
| CCNE1 | Cell cycle | Cyclin E1 binds CDK2 regulating DNA replication and cell cycle progression [113]. |
| CCND1 | Cell cycle | Cyclin D1 forms a complex with CDK4/6 and regulates G1 [113]. |
| TP53 (p53) | TF | Regulates various stress responses including cell cycle arrest, inflammation, autophagy, and apoptosis [114]. |
| FOXO4 | TF | Regulates pathways like oxidative stress signalling, cell cycle progression, and apoptosis [115]. Inhibition acts as a senolytic [116]. |
| FOXO3 | TF | Regulates pathways like oxidative stress signalling, cell cycle progression, and apoptosis [115, 117]. |
| CEBPB (C/EBPβ) | TF | Regulates CS and the SASP [118]. |
| ATF4 | TF | Regulates adaptive genes enabling cells to endure stress like hypoxia or amino acid depletion, but promotes apoptosis under persistent stress [119]. |
| MITF | TF | Promotes proliferation and inhibits apoptosis [120]. |
| EGR2 | TF | Regulates CS, knockdown in RS reverses the senescent phenotype [54]. |

#### 2.1.3 Metabolic heterogeneity

CS is associated with shifts in metabolic profiles [68, 121]. We compiled various metabolic pathways associated with CS from WikiPathways, KEGG, and MetaCyc, alongside a background of metabolic enzymes (see 5.6 Metabolism Pathways in Methods) [122–125]. ORA of metabolic pathways against each other using the metabolism background shows that most pathways comprise unique metabolic enzymes (SI Figure 12, SI Table 18-19).

ORA was performed between metabolism pathways and arrest-DEGs using the intersection of the metabolism background and genes expressed in the fibroblast data as the background (SI Figure 13, SI Table 20). While there was not a significant association between arrest-DEGs and metabolism for most pathways, there was significant upregulation of the eicosanoid metabolism via cyclooxygenases (COX) pathway in SIPS and the hexosamine pathway in OIS, alongside significant downregulation of nucleotide synthesis in contact-inhibited CQ and SIPS. Furthermore, there were various trends, particularly in how energetic pathways – oxidative phosphorylation (OXPHOS), TCA, and glycolysis – were expressed in CQ compared to CS.

We wondered whether samples could be clustered into their respective arrest phenotypes based on metabolic gene expression. While these genes were not capable of clustering RS samples together – or distinguishing between CQ states – hierarchical clustering correctly clustered most samples into CQ, proliferating, SIPS, and OIS groups (SI Figure 14).

Finally, gene expression data was mapped to the eicosanoid metabolism via COX pathway because this pathway is specifically linked to prostaglandins and inflammation [27, 126] and because some of the most variably expressed metabolic genes across all samples were from this pathway, including *PTGS2*, *PTGIS*, and *PTGDS* (Figure 4a, SI Figure 15, SI Table 2). We found that *PLA2G4A* was significantly overexpressed across arrest conditions, except for RS, while *TBXAS1* was uniquely overexpressed in OIS and various metabolic enzymes like *PTGES* and *PTGDS* were shared between SIPS and CQ, but not OIS or RS. In addition, broader trends become apparent even where differential expression did not reach significance individually. In particular, the central enzyme *PTGS2* showed upregulation in SIPS and OIS, no change in RS and serum-inhibited CQ, and downregulation in contact-inhibited CQ; in line with similar patterns for non-metabolic inflammation genes in Figure 4a.

#### 2.1.4 Heterogeneous SASP at the Proteomic Level

Basisty et al. developed the SASP atlas, profiling secretory SASPs (sSASPs) and extracellular vesicle SASPs (eSASPs) from senescent fibroblasts induced via RAS overexpression, irradiation, and atazanavir co-culture, alongside senescent epithelial cells induced via irradiation [49]. The SASP atlas was constructed by comparing secretomes from senescent cells to serum-starved CQ controls, resulting in two groups: i) proteins secreted following CS induction compared to CQ (log_2_(CS/CQ)>0.58 & p-value<0.05 following BH correction); ii) proteins secreted following CQ induction compared to CS (log_2_(CS/CQ)<-0.58 & p-value<0.05 following BH correction) (SI Table 21). The study noted significant heterogeneity in SASP proteins based on stressor and cell type, and distinct secretory profiles between sSASPs and eSASPs.

To compare senescent and quiescent transcriptomes to the SASP atlas, we generated DEGs between CS and CQ samples from the recount3 studies. Limited studies featuring both CS and CQ conditions resulted in a smaller sample size for DEG analysis. Contact-inhibited and serum-starved CQ samples were analysed together against OIS and SIPS samples, excluding RS samples due to insufficient sample numbers (SI Table 1).

After removing the study batch effect via linear regression, PCA was performed whereby samples clustered into three groups corresponding to CQ, SIPS, and OIS samples, and the top two PCAs captured 83% of the sample variance (SI Figure 16). Significant DEGs were generated between CQ and both SIPS and OIS samples, resulting in 3,800 underexpressed and 3,052 overexpressed OIS DEGs, alongside 912 underexpressed and 1,473 overexpressed SIPS DEGs (p<0.05 and |log_2_FC|>log_2_(1.5), negative binomial distribution with BH FDR correction) (SI Table 22).

ORA was performed between SIPS and OIS DEGs generated against CQ samples, with significant overlaps between CS DEGs changing in the same direction using genes expressed in these recount3 samples as the background (SI Figure 17, SI Table 23). ORA was further performed between these DEGs and the stress response pathways; both CS conditions significantly overlapped proinflammatory conditions compared to CQ samples (SI Figure 18, SI Table 24). However, overexpressed OIS DEGs specifically overlapped the p53 pathway, MYC targets, and MTORC1 signalling, whereas SIPS DEGs did not, indicating that fibroblast OIS is specifically associated with these pathways compared to fibroblast SIPS and CQ. Lysosomal genes did not significantly overlap any DEGs, likely because CQ is also significantly associated with lysosomal processes.

ORA was further performed between these DEGs and the SASP atlas by condition and direction using the intersection of genes expressed within the recount3 data and protein secretions from the given SASP condition as the background (Figure 4b, SI Figure 19a, SI Table 25). The overexpressed OIS and SIPS DEGs significantly overlapped the fibroblast RAS sSASP profile, while CQ secretions generated against epithelial and fibroblast irradiated sSASPs were significantly overrepresented amongst downregulated OIS and SIPS DEGs. However, the irradiated fibroblast sSASP and eSASP were only significantly over- and underrepresented in the OIS DEGs, and not the SIPS DEGs. These findings imply that significant portions of some proteomic SASP profiles are captured at the transcriptomic level in a context-dependent manner.

ORA was performed to determine whether SASP profiles are associated with stress response pathways (SI Figure 19b, SI Table 26). None of the SASP profiles were significantly associated with inflammation pathways. However, irradiated SASP secretions specifically were significantly associated with various processes including MTORC1 signalling and hypoxia, whereas other SASP profiles were not, suggesting that SRMs may be partially regulated and effected via the SASP in specific contexts.

### 2.2 Temporal Dynamics of Senescent Cell Transcriptomes

We sought to further dissect the temporal dynamics of CS. Hernandez-Segura et al. generated bulk RNA-seq datasets for fibroblasts, melanocytes, and keratinocytes at 4-, 10-, and 20-days following exposure to 10Gy of γ-radiation, alongside proliferating controls [47] (see 5.1 Cell Cycle Arrest Transcriptomic Data) (SI Table 27). In this work, the researchers identified 61 genes that were shared across all cell types and time points compared to proliferating controls, 34 of which were not shared with quiescent phenotypes.

#### 2.2.1 Heterogeneity of Temporal Senescent States

PCA using the 500 most variable genes showed that CS samples tended to cluster by days post-irradiation, except in the 10-day post-irradiated fibroblasts, which clustered into two groups (SI Figure 20). The keratinocytes had a batch effect that was not present in the other cell type data and was removed via linear regression (see 5.2 Linear Regression in methods) (SI Figure 20a).

Temporal DEGs were generated between each time point and proliferating controls, by cell type, using DESeq2 (p<0.05 and |log_2_(fold change)|>log_2_(1.5), negative binomial distribution with BH correction) [87] (SI Table 28, 29). Unsupervised hierarchical clustering was performed using the top 25 over- and underexpressed temporal DEGs generated between time points compared to proliferating controls (identified using π scores, see 5.3 PCA and Heatmaps) (SI Table 30); samples tended to cluster by days post-irradiation except for one melanocyte sample (SI Figure 21-23).

Temporal DEGs generated between proliferating samples and senescent cells were compared, and significantly more DEGs were shared between all time points for each cell type than expected by chance, based on 10,000 simulations (see 5.4 DEG Overlap Simulations) (SI Table 31-32).

We found four temporal DEGs that were underexpressed across all time points and all cell types – *PIR*, *STMN1*, *USP13,* and *PEG10* – alongside 45 overexpressed temporal DEGs – *AC099489.1, ADM, AL031777.2, AL583836.1, ANKRD29, APLP1, BTG2, C3, CCND1, CNGA3, COLQ, COMP, CSF2RB, DPP6, FGF11, FOLR3, FSTL4, GABBR2, GDNF, H2AC18, H2AC19, H2BC6, H2BC8, H4C8, HES2, IL32, INHA, LIF, LIX1, LTO1, MYOZ2, NECTIN4, PARM1, PLA2G4C, PLXNA3, PTCHD4, PTPRT, RRAD, SERINC4, SIK1, SIK1B, SMCO1, SULF2, WNT9A,* and *ZNF610.* Across all 10,000 DEG overlap simulations, only one DEG was ever shared across all under- and overexpressed time points, indicating that more shared DEGs are significantly conserved across conditions than expected.

The number of shared DEGs across all time points differs from the 61 genes identified in the original study, perhaps because they did not adjust for the keratinocyte batch effect and used a different log_2_(FC) cut-off of log_2_(1.3) instead of log_2_(1.5) [47]. Nonetheless, we rediscovered 42 of the same DEGs.

ORA was performed between cell types by time points, using genes expressed in either cell type as the background (Figure 5a, SI Figure 24, SI Table 33). Various results were as expected. For example, across all cell types and time points, there were more shared overexpressed DEGs than expected by chance (p<0.05, two-tailed Fisher’s exact test with Bonferroni correction). Furthermore, genes that changed in opposite directions between fibroblasts and keratinocytes overlapped significantly less than expected by chance across all time points too. Moreover, DEGs underexpressed in keratinocytes significantly overlapped DEGs underexpressed in fibroblasts across all time points. However, the melanocytes showcased expression patterns opposite to expectation. For example, overexpressed late-senescence (day 20) keratinocyte DEGs overlapped with underexpressed melanocyte DEGs more than expected by chance across all time points. Furthermore, overexpressed melanocyte DEGs significantly overlapped the underexpressed keratinocyte and fibroblast DEGs across most time points. These results suggest that the irradiation-induced CS programme in melanocytes is distinct from keratinocytes and fibroblasts.

**Figure 5.**
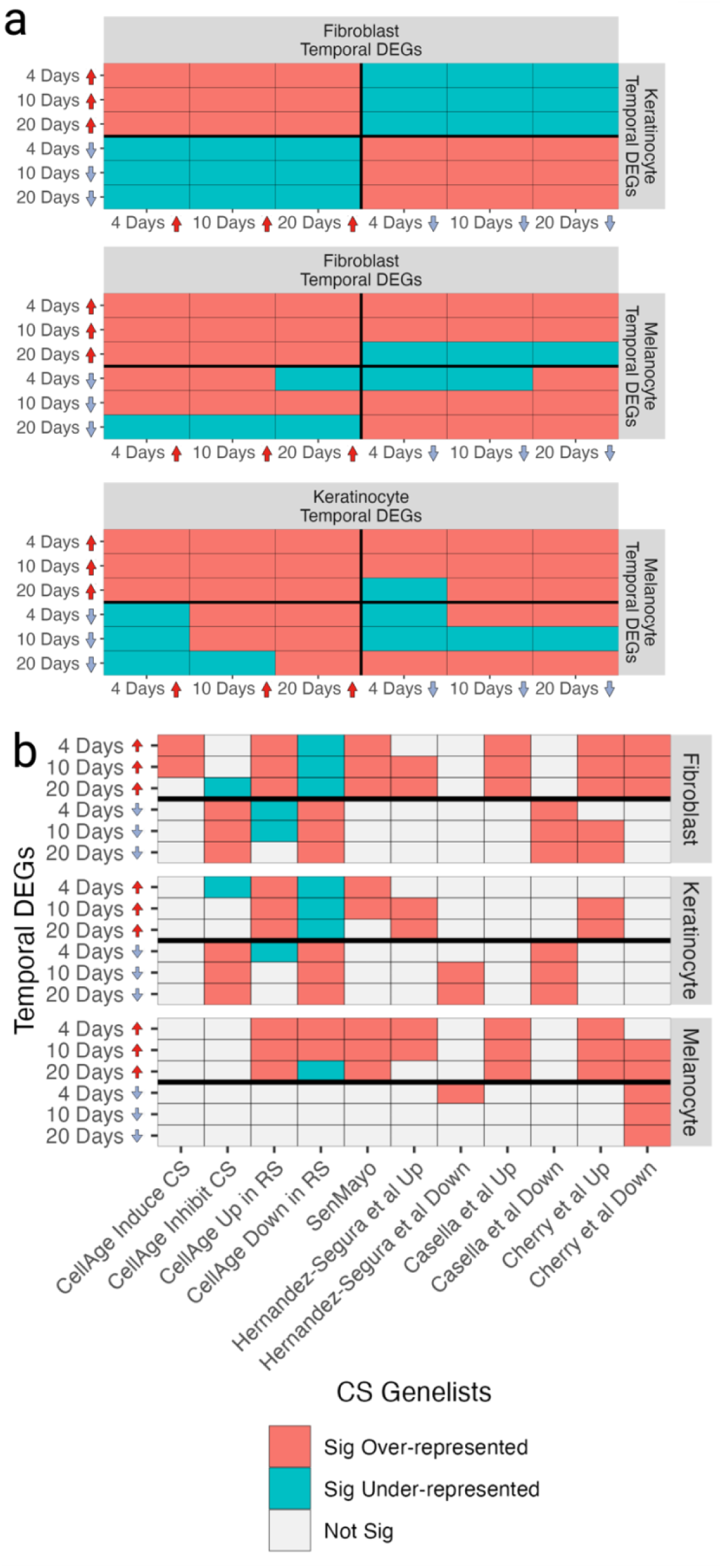
a) Overlap between temporal DEGs. b) Overlap of temporal DEGs with various gene lists of interest.

ORA was performed between temporal DEGs and the CS datasets, using genes expressed in each cell type as the background (Figure 5b, SI Figure 25, SI Table 34). Overexpressed CellAge signatures of RS were significantly upregulated across all time points, and underexpressed CellAge signatures were downregulated in fibroblasts and keratinocytes (p<0.05, two-tailed Fisher’s exact test with Bonferroni correction). However, the RS signatures were significantly upregulated in early- and mid-melanocyte time points. Moreover, CellAge inhibitors of CS were significantly downregulated in fibroblasts and keratinocytes, but not melanocytes, further highlighting differences in the melanocyte irradiation-induced CS programme. Hernandez-Segura et al. signatures overlapped the temporal DEGs as expected, which is not surprising given these signatures were partially derived from the 10-day data, although not all overlaps were significant. The closest universal signature of CS was SenMayo, which was consistently upregulated across cell types and time points, although late-stage keratinocytes did not significantly upregulate SenMayo signatures, indicating a false negative. Furthermore, none of the 10 SenMayo DEGs we identified as universally overexpressed in arrest-DEGs (SI Figure 8) were significantly overexpressed across all cell types and time points, further suggesting that SenMayo is measuring heterogeneous biological processes.

#### 2.2.2 Temporal Activation of Stress Response Genes in Arrest Phenotypes

ORA was performed between temporal DEGs and the aforementioned pathways using genes expressed in the temporal samples as the background (Figure 6a, SI Figure 26, SI Table 35). There was significant heterogeneity among the temporal DEGs. The only uniformly significantly overexpressed pathway across cell types and time points was the ‘TNFA signalling via NF-κB’ pathway, with the p53 pathway being significantly overexpressed across time points except in early keratinocyte CS. Lysosomal DEGs and MYC targets were only significantly over- and underexpressed in fibroblasts, respectively. Inflammatory pathways were heterogeneously expressed on a cell type and temporal basis, alongside the hypoxia pathway. While melanocytes and fibroblasts significantly upregulated apoptosis pathways across time points, this was not observed in keratinocytes. Notably, melanocytes did not downregulate any pathways except for MYC targets at late CS. This includes various proliferative pathways — mitotic spindle, G2M checkpoint, and E2F targets — that were otherwise downregulated in fibroblasts and keratinocytes as expected. At early- and mid-CS time points, melanocytes significantly upregulated E2F targets, contrary to expectation, suggesting that these cells may maintain proliferative capacity despite upregulating other stress response pathways.

**Figure 6.**
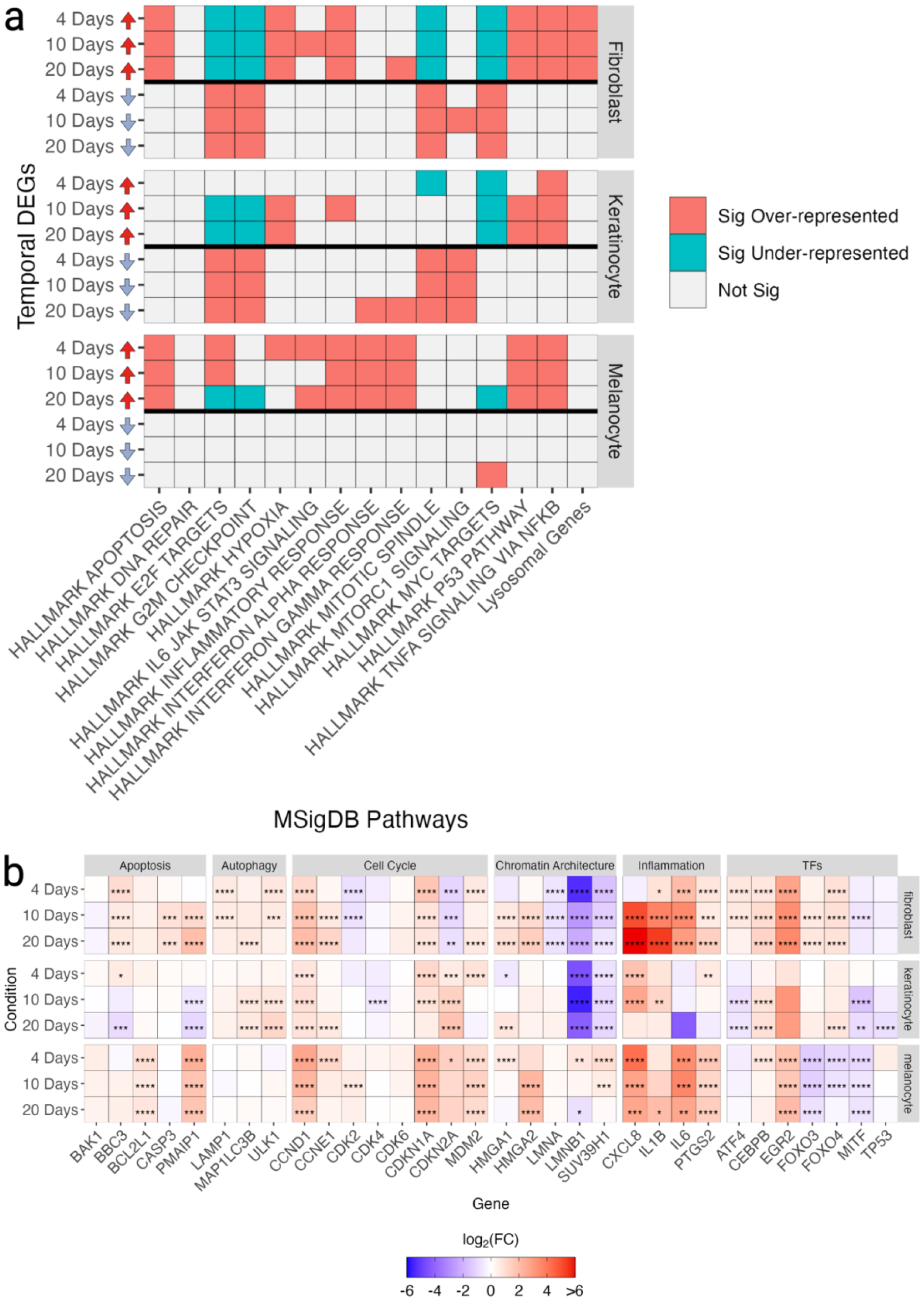
a) Overlap between temporal DEGs and various pathways of interest. b) Expression of key stress response genes across irradiated cell types by days post-irradiation, compared to proliferating controls.

The expression of various stress-response and senescence-associated genes across irradiated cell types and time points was assessed (Figure 6b). Hernandez-Segura et al. previously showed that various driver genes of CS like p21, p16, and p53 are not differentially expressed across all time points [47]. Indeed, we found a lack of universal expression of *CDKN1A*, *TP53*, and *CDKN2A* within this data. Furthermore, the aforementioned stress response genes were expressed heterogeneously, dependent on cell type and time point. Moreover, there were differences in key TFs and chromatin architecture genes depending on context. A particularly interesting example is *LMNB1*, which was significantly downregulated across time points in the fibroblast and keratinocyte temporal data but was upregulated in early melanocyte CS and only significantly downregulated at late melanocyte CS. Intriguingly, we also found keratinocytes to be less proinflammatory than melanocytes or fibroblasts, with *IL6* never overexpressing in any keratinocyte time point.

## 3. Discussion

### 3.1 An Undefinable Phenotype

The concepts of ‘senescence’ and ‘quiescence’ as heterogeneous cell cycle arrest phenotypes are well-established [47, 72, 127–129]. However, the current understanding of senescence is riddled with paradoxes and contradictions. Classical markers of senescence include the absence of cell cycling (e.g., lack of Ki-67), expression of cyclin-dependent kinase inhibitors (e.g., p53/p21), secretion of proinflammatory factors (e.g., IL6), and increased lysosomal activity (e.g., β-gal staining) [22]. Yet, these markers are also found in other contexts and have all been uncoupled from ‘senescent’ states (Table 1).

We argue that mosaic co-activation of clusters of genes that modulate distinct stress responses (Stress Response Modules, SRMs) encompass what the field refers to as ‘heterogeneous senescence phenotypes.’ Under this model, the aforementioned markers of senescence represent markers of distinct SRMs (Figure 1). Because SRMs are heterogeneously expressed and no single SRM is universally guaranteed to be expressed across all biological contexts, the result is a biological phenomenon that cannot rigorously be defined – ‘senescence.’ Here, we discuss implications of this model, while SI Document 1 provides supporting evidence from our results for some of the points we discuss.

### 3.2 Variable Stress Response Pathways Across Arrest State

Our results indicate that arrest transcriptomes are associated with various stress response pathways including lysosomal activity, inflammation, apoptosis, and hypoxia, in a context-dependent manner (SI Document 1). This is evident at both the pathway level (Figure 3a, 6a, and SI Figure 18) and in the expression of key genes linked to senescence- and stress-associated phenotypes (Figure 4a and 6b).

### 3.3 SRM Regulation and Crosstalk

CS literature suggests that various processes regulate different aspects of arrest phenotypes. Here, we discuss mechanisms potentially underlying how SRMs are controlled at the transcriptomic, chromatin accessibility, and metabolic level.

#### 3.3.1 Transcription Factors

Various key regulators of stress response pathways, including *TP53* and its regulator *MDM2*, alongside *ATF4* – the main effector of the integrated stress response (ISR) – were differentially expressed in a context-dependent manner (Figure 4a and 6b, SI Document 1). This was also the case for *CEBPB*. Importantly, these genes can regulate various stress responses, including inflammation, autophagy, and the SASP, suggesting that heterogeneity amongst TFs could play a role in modulating distinct SRMs amongst heterogeneous ‘senescent’ cell populations [64, 114, 130–134].

#### 3.3.2 Chromatin Rewiring

Chromatin accessibility regulates SRMs (SI Document 1). For example, *IL1B* – a proinflammatory SASP factor which upregulates in various senescence phenotypes (Figure 4a and 6b) [27] – is also upregulated in TNFα-treated cells, alongside SASP factor *IL1A* and cell cycle gene *CKAP2L* [65]. In OIS, the upregulation of these three genes involves global epigenetic alterations in chromatin accessibility, resulting in enhancer-promoter rewiring [65]. While these three genes are also upregulated in TNFα-treated cells, this process is mediated via TFs and not chromatin rewiring [135]. Furthermore, senescence-associated cell cycle arrest is reversible in some contexts – such as via p53 knockdown in fibroblasts, provided p16 expression remains low [39]. While these cells maintain their proliferative capacity, SASP factors continue to be secreted [27, 64]. These studies suggest that chromatin rewiring plays a role in determining the activation and reversibility of SRM activation in CS and other contexts [27, 136].

#### 3.3.3 Heterogeneous Cellular Signalling

We found that irradiation-induced SASPs specifically are linked to angiogenesis, coagulation, hypoxia, and MTORC1 signalling, indicating potential partial regulation of some SRMs by the SASP under specific circumstances (Figure 5b). Senescent cells can induce CS in a paracrine and juxtacrine manner, and reinforce ‘senescent’ states via autocrine signalling [137–140]. Paracrine and juxtacrine signalling, which mediate secondary senescence, has likely evolved in part to amplify wound healing and immune system signalling [141]. Nonetheless, cells entering secondary senescence are distinct from primarily senescent cells [139, 140], as they must be in order to inhibit the uncontrolled propagation of CS states [142], indicating that SASP-induced SRM regulation and/or execution is distinct to primarily ‘senescent’ cells, and context-dependent.

#### 3.3.4 Metabolism

Metabolism is implicated in regulating SRMs and various metabolic pathways are implicated in CS (SI Table 36) [121, 143, 144]. From the patterns and trends observed in our arrest-DEG ORAs (Section 2.1.3), multiple connections to other features of CS and CQ become apparent. Nucleotide synthesis is universally downregulated in all arrest conditions, while energy metabolism is sustained in CS but not CQ. Inflammation-related metabolic pathways agree with other CS type-specific inflammation features, as does NAMPT in NAD salvage (see SI Document 1 for more details).

Overall, the targeted analysis of metabolic pathways hints at relevant connections to non-metabolic aspects of CS and CQ, with condition-specific differences. Furthermore, despite the subtle nature of the metabolic pathway alterations, they were significant and consistent enough to cluster arrested and proliferating samples into their respective categories, excluding RS (SI Figure 14). This indicates that the differences in metabolic profiles are distinct and reliable enough to categorise cells based on the specific insult used to arrest them. Going forward, the study of metabolic alterations in CS and CQ faces specific challenges, suggesting that to further investigate the metabolic features of senescence, systems biology methods that study network functionality beyond pathway-level statistical measures might be advantageous [121].

#### 3.3.5 Crosstalk between SRMs

Senescence has been considered a ‘heterogeneous’ phenotype as opposed to differential activation of SRMs because SRMs are often co-activated, likely via multiple mechanisms. The most obvious case is that SRMs are sometimes regulated via the same mechanism. For example, RAS-induced CS triggers chromatin rewiring which facilitates enhancer-promoter interactions linked to both upregulation of inflammation and downregulation of cell cycle genes [65]. Furthermore, the MSigDB apoptosis pathway is significantly associated with hypoxia, inflammation, and MTORC1 signalling, indicating crosstalk between SRMs (SI Figure 11). Indeed, transition into senescent-like states involves positive feedback loops that reinforce DDR signalling via SASP factors, chromatin remodelling and degradation, and mitochondrial dysfunction and ROS [145].

Nonetheless, it is also possible that dysregulating one SRM causes internal stress within the cell, resulting in the activation of other SRMs. CellAge genes – which directly induce or inhibit ‘senescence’ when manipulated genetically [21, 88] – are significantly associated with various pathways, including apoptosis and MTORC1 signalling (Figure 3b). Furthermore, autophagy and CS have a complex relationship, and both induction and inhibition of autophagy promote CS [146–148]. More research is needed to understand the logic that underlies SRM regulation.

### 3.4 Implications of CS as Heterogeneous Activation of SRMs

Redefining CS from a heterogeneous phenotype to the heterogeneous activation of stress responses may seem like a semantic argument. However, we argue that this proposed paradigm shift offers explanations for various paradoxes in mammalian biology. Our model provides explanations for the various idiosyncrasies within the CS field (Table 1) and has implications for ageing, cancer, and chronic diseases [149, 150].

#### 3.4.1 Lack of a Universal Signature

We identified that various fibroblast DEGs induced into RS, SIPS, and OIS are significantly shared across arrest phenotypes – including CQ. Previous studies have also identified and published transcriptomic signatures and markers associated with cellular senescence (Table 2). This prompted us to explore whether these gene lists could serve as universal signatures of CS, based on the following criteria: i) being unique to CS and not other biological phenomena; and ii) being universally differentially expressed across all CS conditions. However, none of the CS gene lists meet these criteria (Figure 2b and 5b). For example, SenMayo is a strong contender, significantly and exclusively enriching across most CS conditions, except for keratinocytes sequenced 20 days post-irradiation. However, only 10 genes were overexpressed across all CS DEGs (SI Figure 8, SI Table 12), and none of these genes are shared among the universally overexpressed temporal DEGs. Moreover, some of these CS gene lists – including SenMayo – also significantly enrich for various stress responses, indicating they may be measuring distinct SRMs as opposed to universal senescence-specific processes (Figure 3b). Indeed, activated macrophages enriched for the SenMayo gene list as well (SI Figure 10), and SenMayo itself is known to be over-reliant on proinflammatory SASP factors [23].

#### 3.4.2 Temporal Dynamics of CS

Previous studies have shown a sequential order of gain-of-senescence phenotypes in cells that enter CS [61, 63]. As has been discussed, various genes linked to inflammation, autophagy, and apoptosis show dynamic gene expression over time (Figure 6b). Under our model, previously reported shifts in senescence phenotypes at least partially represent the temporal regulation of SRMs.

#### 3.4.3 Post-Mitotic CS

Post-mitotic cells like neurons and skeletal muscle cells are reported to activate senescent states under stress, assessed via p16 and p21 expression, secretion of a SASP, and β-gal staining [52, 151–153]. However, these cells – being terminally differentiated — do not proliferate to begin with. Under our model, these examples constitute cells that have activated autophagy- and inflammation-associated SRMs in response to stress, independent of cell cycle arrest.

#### 3.4.4 CS in Cancer

Senescent cancer cells are associated with relapse and treatment resistance in various instances, including acute myeloid leukaemia and triple-negative breast cancer [154, 155]. While cell cycle arrest is a barrier to cancer formation, cancer cells are known to hijack senescence pathways to drive tumorigenesis and promote survival. Indeed, inflammation and autophagy can be manipulated to encourage tumorigenesis and treatment resistance and inhibit apoptosis [27, 156, 157]. Furthermore, studies show that cancer cells can upregulate anti-apoptotic pathways associated with senescence induction like BCL-2 and BCL-XL [158, 159].

An important point to consider is that multiple mechanisms of cell cycle arrest may have evolved in mammals (Figure 1). Importantly, the p53/p21 pathway is also associated with reversible arrest [35–38], whereas p16 appears to yield less reversible forms of CS [64]. While studies purport to have reversed RS via p16 knockdown [53, 54], a strong contender for less-reversible forms of ‘senescence’ are chromatin rearrangements and perhaps formation of SAHF, which are more strongly associated with OIS, not RS [32, 100]. Our model of SRM induction sheds light on potential strategies that cancer cells might utilise to co-opt CS and CQ SRMs to enhance their survival, perhaps by hijacking the reversible form of cell cycle arrest in response to cancer treatments instead of activating irreversible arrest [160].

#### 3.4.5 CS in Ageing and Chronic Diseases

Various stress responses dysregulate with age, including an increase in inflammation and a decrease in autophagy, alongside the accumulation of ‘senescent’ cells [161–163]. Eliminating senescent cells extends the lifespan and healthspan of mice [164, 165], and research utilising mouse models has indicated that senolytics can lead to improvements in the pathological features of various ageing-related diseases, including diabetes [166], Alzheimer’s disease [167], and osteoarthritis [168, 169]. The implication is that the accumulation of cells with dysregulated SRMs is potentially driving – at least partially – ageing and ageing-related pathologies, and the elimination of cells with dysregulated SRMs has positive outcomes. More research is necessary to determine what drives the dysregulation of SRMs with age and disease, although, given the multifactorial nature of these phenotypes, there are likely various sources that lead to the dysregulation of SRMs with age.

Senolytics currently suffer from two drawbacks: i) off-target effects; and ii) senolytic-resistant ‘senescent’ cells [170, 171]. Finding distinct drugs to selectively target cells with specific and distinct dysregulated SRMs – stressolytics – could be a potential future avenue for understanding and treating pathologies associated with the accumulation of cells with specific dysregulated SRMs. Senomorphics capable of modulating specific and distinct SRMs may also be applicable [172].

#### 3.4.6 CS in Benign Contexts

There is evidence of programmed ‘senescence’ in embryogenesis, while senescent cells play a role in normal physiology including wound healing, tissue repair, and embryo receptivity [173–175]. In the context of mammalian development, CS and apoptosis are programmed processes involved in limb formation, and aberrant regulation leads to developmental defects [174, 176]. Under our model, these cells have evolved to activate SRMs to achieve beneficial outcomes, including autophagy and inflammation in wound healing [177]. We acknowledge that SRMs may be activated in contexts independent of biological ‘stress’ – like contact-inhibited quiescence. Nonetheless, these same pathways appear to also activate across stress phenotypes – dependent on context – and the term ‘stress response module’ is therefore apt.

### 3.5 Future Direction

While this new perspective is promising, further research is needed to specifically define SRMs, and to identify how they are regulated and how they modulate different phenotypes including senescence-associated phenomena. We have been hesitant to clearly define SRMs because it is unclear to what extent processes like autophagy and the SASP can be subdivided into more specific SRMs themselves, such as mitophagy or ECM remodelers.

Rather than focusing on a universal marker of senescence, there should be a focus on finding robust markers for individual SRMs. Single-cell RNA-seq of cells under various stress conditions will further allow for the identification of SRMs. Focusing on the dynamics of SRMs – as opposed to a ‘heterogeneous’ senescent phenotype – will clarify the role of variable stress responses in ageing, development, and disease, amongst other biological phenomena.

## 4. Conclusion

We demonstrated that senescent cells exhibit heterogeneous transcriptomic and secreted proteomic changes associated with diverse stress response pathways, including inflammation, autophagy, and apoptosis, in a cell-type, temporal, and insult-dependent manner. CS signatures reported in the literature are inadequate for exclusively identifying senescent cells across all contexts, emphasising the need for a more nuanced approach. We propose that ‘senescent’ cells lack a universal marker because senescence is a mosaic differential activation of various stress-associated pathways – with distinct phenotypic biomarkers – dependent on context. We call the clusters of genes that control and effect these stress responses Stress Response Modules (SRMs), and propose that no consistent ‘core’ of SRMs exists that could be used to meaningfully define the senescent state.

We find that TFs, genes controlling chromatin accessibility, and metabolic enzymes are heterogeneously expressed in a context-dependent manner. Additionally, some SASP profiles – which can be partially identified at the transcriptomic level – also enrich for various stress responses, indicating multiple avenues by which SRMs are regulated. Our model provides a framework for understanding the role of stress responses across a range of biological contexts, while also exploring the regulatory mechanisms underlying senescence.

Future research should focus on validating the SRM model through approaches like single-cell RNA sequencing to determine the logic underlying SRM activity. Additionally, understanding how SRMs dysregulate with age and disease, and their role in normal physiology, will lead to novel insights into various biological phenomena.

## 5. Methods

### 5.1 Cell Cycle Arrest Transcriptomic Data

Lung, skin, and foreskin fibroblast CQ and CS data were obtained by manually annotating and filtering recount3 for relevant studies and downloaded using the recount3 R package [85, 86]. We only included samples that were not under multiple treatments (e.g., we only included cells that were irradiated or starved, not both). Samples were included if they were clearly labelled as proliferative, senescent, or quiescent, alongside the mechanism which was used to induce cell cycle arrest. We excluded samples with genetic manipulations unless the manipulation was neutral, such as scramble siRNAs or GFP inserts, or if the genetic manipulation was used to induce OIS (i.e., HRAS or BRAF overexpression). The following studies were excluded as they added noise to the PCA plots and did not cluster well: SRP050179, SRP195418, SRP113334, SRP127595.

Samples were downloaded using the recount3 R package via the *create_rse* function [85, 86]. Sample numbers are available in Table 6. Because the data was derived from various studies, counts were scaled using the *transform_counts* function from the recount3 R package. Samples were selected if they met the following criteria:

- The samples comprised bulk RNA-seq data from non-transformed human lung, skin, or foreskin fibroblasts. Both primary cells and cell lines were included.
- For proliferating, RS, OIS, serum-starved and contact-inhibited CQ, samples were included if the authors of the study labelled the cells as such. We could not find suitable heat shock CQ samples.
- For SIPS, we included samples that were induced into DNA-damage induced CS via co-culture with bleomycin (n=7), etoposide (n=8) or hydrogen peroxide (n=3), or irradiated with 10Gy (n=21).

Temporal irradiation-induced CS data and proliferating controls for fibroblasts, keratinocytes, and melanocytes were obtained from Hernandez-Segura et al. (ArrayExpress accession E-MTAB-5403) [47]. For each cell type and condition, there were 6 samples.

**Table 6.** Number of arrested and proliferating fibroblast samples that were included from recount3, by tissue type.

| Tissue | Cell State | Sample n |
| --- | --- | --- |
| Foreskin | Proliferating | 25 |
| Lung | Proliferating | 61 |
| Skin | Proliferating | 5 |
| Foreskin | Contact-inhibited CQ | 8 |
| Lung | Contact-inhibited CQ | 3 |
| Skin | Contact-inhibited CQ | 8 |
| Foreskin | Serum-starved CQ | 13 |
| Lung | Serum-starved CQ | 9 |
| Skin | Serum-starved CQ | 0 |
| Foreskin | Replicative CS | 2 |
| Lung | Replicative CS | 9 |
| Skin | Replicative CS | 0 |
| Foreskin | Stress-induced CS | 18 |
| Lung | Stress-induced CS | 18 |
| Skin | Stress-induced CS | 3 |
| Foreskin | Oncogene-induced CS | 7 |
| Lung | Oncogene-induced CS | 41 |
| Skin | Oncogene-induced CS | 0 |

Genes with more than 1 count per million in at least 30% of samples for any given arrest condition were included for the analyses, and we limited our analysis to protein-coding genes, downloaded using biomart version 100 via the biomaRt R package [178–181].

### 5.2 Linear Regression

We found DEGs for each cell type between the various time points post-irradiation compared to proliferating controls in the time-series data, alongside DEGs between arrested and proliferating cells in the recount3 data. DEGs were also generated between SIPS and OIS samples compared to grouped serum-starved and contact-inhibited CQ samples. DEGs were generated between arrest conditions, as opposed to other variables like tissue type. Linear regression was used to account for batch effects within the data. For lung, skin, and foreskin fibroblasts induced into cell cycle arrest using various insults, the following regression model was used, with the total number of DEGs outlined in Table 7:

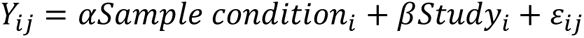

**Table 7.** Number of significant DEGs by condition, compared to corresponding proliferating controls.

| Group | Direction vs Proliferating | DEG n |
| --- | --- | --- |
| Contact-inhibited CQ | Down | 2,110 |
| Serum-starved CQ | Down | 1,565 |
| RS | Down | 1,966 |
| SIPS | Down | 883 |
| OIS | Down | 2,496 |
| Contact-inhibited CQ | Up | 3,457 |
| Serum-starved CQ | Up | 2,698 |
| RS | Up | 1,809 |
| SIPS | Up | 1,784 |
| OIS | Up | 1,689 |

For OIS and SIPS DEGs generated against CQ DEGs, the following regression was used:

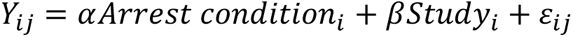

Importantly, contact-inhibited and serum-starved CQ samples were grouped to increase sample size, and DEGs were generated between individual CS conditions against the CQ samples.

For melanocyte and fibroblast temporal data, the following regression model was used:

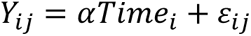

The keratinocyte data had an obvious batch effect (SI Figure 20a). We contacted the study’s corresponding author, Marco Demaria, and it appears that the keratinocyte data was processed by two researchers. As such, we opted to manually label this batch effect, and account for it using the following regression model:

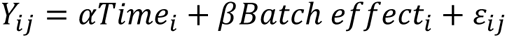

Variables were defined as follows:

- Yij: The expression level of gene j in sample i.
- Sample condition: OIS, SIPS, RS, contact-inhibited CQ, serum-starved CQ, or proliferative state of each sample.
- Arrest condition: OIS, SIPS, or grouped CQ state of each sample.
- Time: The number of days following exposure to 10Gy ionising radiation.
- Batch effect: The manually labelled batch effect within the keratinocyte data.
- εij: The error term for gene j in sample i.

The *DESeq* and *results* functions from the DESeq2 R package v1.36.0 were used with default parameters to generate DEGs [182, 183]. The *results* function has an independent filtering option which was used for higher statistical power to obtain more biologically meaningful results, as specified in the DESeq2 documentation [184]. The *results* function also provides Cook’s distances, which were used to remove outliers [185]. DEGs were considered significant if they had an adjusted p-value<0.05 (negative binomial distribution with BH correction) and |log_2_(fold-change)|>log_2_(1.5). Volcano plots were generated using the *EnhancedVolcano* function from the EnhancedVolcano R package [186].

### 5.3 PCA and Heatmaps

Blinded variance stabilising transformations (VST) were performed on the data prior to PCA using the *varianceStabilizingTransformation* function from the DESeq2 R package with default parameters [87]. PCs were calculated using the top 500 most variable genes and plotted using the *plotPCA* function.

Heatmaps with hierarchical clustering were generated using the *pheatmap* function from the pheatmap R package [187]. Briefly, we applied a blinded VST normalisation to all the counts data using the *varianceStabilizingTransformation* function from the DESeq2 R package [87], and then filtered the counts data for DEGs before running *pheatmap*; normalised gene count were scaled using the scale=’row’ argument. Only the top DEGs were used to generate heatmaps, with the exact number of DEGs used specified after each heatmap. We calculated a π score for each DEG using the following equation [188]:

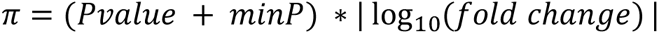

Where Pvalue was the adjusted p-value for each gene, minP was the minimum p-value for any set of DEGs (to avoid multiplying by 0 when the calculated p-value was below the minimum floating-point number allowed by R), and fold change was the fold change for the respective DEG. We then extracted the top genes based on π scores and used these to plot heat maps [188]. For the metabolism heatmap, all genes were used to cluster samples and as such a π score was not calculated.

For plotting both PCA plots and heatmaps, the batch effects outlined in 5.2 Linear Regression were removed from the counts data using the *empiricalBayesLM* function from the WGCNA package [189], while relevant groups like ‘sample condition’ for arrest-DEGs, ‘arrest condition’ for OIS and SIPS DEGs, and ‘time’ for temporal DEGs were preserved. For calculating gene expression variance across samples, the *rowVars* R function was used on normalised counts.

### 5.4 DEG Overlap Simulations

Simulations can be used to determine the probability of multiple DEGs differentially expressing in the same direction across time points or arrest states. The basic steps are as follows:

1. Find DEGs between conditions. Separate all genes — regardless of whether they are significantly differentially expressed — into over- and underexpressed genes compared to proliferating controls, based on the sign of the fold change.
2. Count the number of DEGs that are significantly over- and underexpressed in each arrested condition.
3. From the pool of overexpressed genes, randomly sample the number of significantly overexpressed DEGs. Repeat the process for each arrested cell state.
4. Overlap the randomly sampled overexpressed genes between all arrested cell states.
5. Repeat steps 3-4 for the underexpressed DEGs.
6. Repeat the sampling process 10,000 times for the overexpressed and underexpressed DEGs.
7. Calculate a probability distribution to determine how many DEGs would be expected to change in the same direction by chance across multiple conditions if the DEGs were completely random.

In total, three simulation instances were performed:

1. Simulation between arrest-DEGs generated from serum-starved and contact-inhibited CQ lung, skin, and foreskin fibroblast samples, alongside RS, OIS, and SIPS samples vs. proliferating controls.
2. Simulation between temporal DEGs 4-, 10-, and 20-days post-irradiation vs. proliferating controls by cell type.
3. Simulations between DEGs shared across all nine temporal conditions (4-, 10-, and 20-day post-irradiated DEGs in fibroblasts, keratinocytes, and melanocytes generated against proliferating controls).

### 5.5 Gene Overlaps

To test for overrepresentation between gene lists, the GeneOverlap R package v1.36.0 was used [190]. Significance between gene lists was assessed using a two-tailed Fisher’s exact test with Bonferroni correction, and the background for each overlap is stated alongside each overlap. Upset plots were generated using the ComplexUpset R package v1.3.3 [191]. For the SASP atlas, secretions were considered significant when BH-adjusted p-values were <=0.05 and the |log_2_(ratio)| of CS to CQ secretions was > log_2_(0.58) or < -log_2_(0.58) for senescent and quiescent secretions respectively, as stated in the original paper [49]. Protein-coding genes were filtered for genes in ensembl version 100 for consistency. Furthermore, when multiple proteins were listed with just one p-value (e.g. CXCL1;CXCL2;CXCL3) these entries were removed to reduce ambiguity for gene overlaps.

For stress response overlaps, we used the MSigDB, which constitutes refined gene sets that convey specific biological states and processes and provides more refined and concise inputs for enrichment analyses [192]. The MSigDB contains two MYC gene lists, which were merged for simplification. Lysosome and lysosome-related genes were also included, derived from a database of genes related to autophagy [93]. While gene overlaps and FDR correction was performed for all MSigDB pathways, we have only plotted the most interesting pathways due to space constraints.

### 5.6 Metabolic Pathways

#### 5.6.1 Building Metabolic Pathways

ORA was performed using a manually curated list of metabolic pathways known to be perturbed in CS (SI Table 18, 36). Entries for these pathways were sourced from the WikiPathways, KEGG, and MetaCyc databases [122–124]. When biologically meaningful, gene lists from related pathways were merged, or subsets of genes were extracted if a pathway entry encompassed multiple subsystems. SI Table 36 lists all pathways, along with the databases and entries their gene lists were based on, as well as any modifications, such as the use of specific gene subsets. For example, non-metabolic genes like TFs were manually filtered from the pathway gene lists (SI Table 37). Furthermore, there are references showing how the given metabolic pathway links to CS.

Given the metabolic gene list was focused around metabolic enzymes specifically, we limited the ORA background to only include metabolic enzymes. In particular, the genes included in the human genome-scale metabolic model Human1 (version 1.17.0) were used as a basis [125]. To form the complete background, the Human1 gene list was merged with the lists of metabolic genes of each pathway and duplicates were pruned (SI Table 18). The constructed complete background thus provides an approximation of the human metabolic genome comprising 2,894 genes, including all chosen pathways of interest.

#### 5.6.2 Mapping Metabolic Pathways

The Cytoscape software was used to visualise the Eicosanoid metabolism via cyclooxygenases WikiPathway with accession WP4347, and map arrest FC data onto it, using the WikiPathways app [193, 194].

## Supporting information

Supplemental discussion text

## Authors’ contributions

RAA wrote the manuscript. RAA annotated recount3 samples. RAA, CL, NK, and MB performed bioinformatics analyses. RAA, CL, and MB interpreted the data. TD, CL, MB, and JPM edited the manuscript. RAA, TD, and JPM conceived the project.

## Funding

CellAge was supported by a grant from the Biotechnology and Biological Sciences Research Council (BB/R014949/1) to JPM. Work in our lab is further supported by grants from the Wellcome Trust (208375/Z/17/Z), Longevity Impetus Grants, LongeCity and the Biotechnology and Biological Sciences Research Council (BB/V010123/1).

## Data Availability

Senescence and quiescence samples were downloaded from recount3 using the following accessions: SRP089801, SRP065206, SRP052706, SRP154382, SRP096629, SRP066947, ERP021140, SRP153205, SRP017378, SRP153724, SRP154577, SRP045867, SRP060598, SRP040243, SRP172671, SRP136071, SRP136727, SRP069768, SRP113329, SRP046254, SRP127037, SRP062872, SRP113324, SRP034163, SRP017142, SRP064207, SRP040745, SRP098713, SRP066917, SRP034541, SRP070636, SRP121031, SRP123346, and SRP117883. Temporal data was downloaded using the following accession: ERP021140. CellAge is available from the HAGR website at https://genomics.senescence.info/cells/ [88]. All code available at the following GitHub repository: https://github.com/avelar-ageing/senescence_stress.

## Competing interests

JPM is CSO of YouthBio Therapeutics, an advisor/consultant for the Longevity Vision Fund, 199 Biotechnologies, and NOVOS, and the founder of Magellan Science Ltd, a company providing consulting services in longevity science.

## Acknowledgements

Thanks to Professor Marco Demaria and Professor Anne McArdle for a fruitful discussion on stress responses in CS and CQ, alongside the Narita lab for providing knowledge on chromatin dynamics in CS, Birgit Veldman for her literature research into the role of metabolic pathways in senescence, and Daniel Palmer for proof-reading the manuscript.

## Notes

https://github.com/avelar-ageing/senescence_stress

