## Supplemental discussion text for "Mosaic Regulation of Stress Pathways Underlies Senescent Cell Heterogeneity"

While the majority of the discussion is focused on outlining implications of SRMs underlying senescent phenotypes, here we provide evidence for some of the discussion points based on our results.

### 3.2.1 Cell Cycle Arrest

There is heterogeneous expression of various cell cycle genes. For instance, irradiation-induced CS results in significant under- and overexpression of *CDKN2A* in fibroblasts and keratinocytes, respectively (Figure 6b). A key finding is that melanocytes upregulate many transcripts that are downregulated in keratinocytes and fibroblasts after identical radiation exposure (Figure 6a). Unlike arrest-DEGs and temporal DEGs from irradiated keratinocytes and fibroblasts, melanocytes do not significantly downregulate proliferation-associated pathways related to the G2M checkpoint, mitotic spindle, and E2F target genes across timepoints (Figure 3a and 6a). Notably, at 4- and 10-days post-irradiation, there is significant upregulation of E2F target genes in melanocytes, indicating that heterogeneity among melanocyte-specific DEGs is partially driven via differences in cell cycle processes. Scientific literature suggests that melanocytes proliferate upon irradiation [1, 2]. Nonetheless, these cells also upregulate other CS-associated stress response pathways, including inflammation, apoptosis, and p53 signalling, and were considered senescent in the original study [3], indicating potential activation of various SRMs independent of cell cycle arrest. However, there may be distinct populations of melanocytes that are either cycling or arrested, which could be masked by bulk RNA sequencing. Single-cell RNA sequencing of irradiated melanocytes will be fundamental for revealing potential heterogeneity within these cells.

### 3.2.2 Autophagy

Across both senescent and quiescent fibroblasts, there appears to be upregulation of lysosomal-associated genes, regardless of insult and temporal dynamics (Figure 3a and 6a). However, keratinocytes and melanocytes induced into CS via irradiation did not upregulate lysosomal transcripts. Nonetheless, various genes important for autophagy and lysosome formation – including *ULK1* and various FOXO TFs [4-6] – were heterogeneously differentially expressed, dependent on context (Figure 4a and 6b). Furthermore, the mTOR pathway – which is a master regulator of autophagy [7] – was heterogeneously dysregulated in an insult- and cell type-dependent manner. For example, MTORC1 signalling was significantly up- and downregulated in OIS and SIPS compared to CQ, respectively (SI Figure 18). Moreover, this pathway was universally downregulated across all irradiated keratinocytes timepoints but none of the melanocyte timepoints (Figure 6a).

### 3.2.3 Inflammation and the SASP

The SASP – perhaps the most important feature of senescent cells – is implicated in wound healing, cancer progression, inflammation, extracellular matrix remodelling, and cellular reprogramming [8-11]. SASPs comprise heterogenous secretions dependent on insult and cell type [12]. Nonetheless, a significant number of SASP proteins are detectable via transcriptomics in a context-dependent manner, particularly for irradiated SASP profiles (Figure 4b). This was the case even when the SASP atlas insult differed from the insult used to generate OIS and SIPS DEGs against CQ samples.

The SASP constitutes various proinflammatory cytokines and chemokines [10]. Intriguingly, fibroblasts induced into RS did not appear to upregulate inflammation-associated genes including *IL6*, *IL1A*, *IL1B*, and *CXCL8*, unlike SIPS and OIS fibroblasts (Figure 4a). Nonetheless, all senescence-associated transcriptomes significantly enriched for various proinflammatory pathways, although the specifics of which inflammation pathways were enriched was context-dependent (Figure 3a and 6a). Furthermore, the temporal DEGs suggest that irradiated fibroblasts are significantly more pro-inflammatory than keratinocytes or melanocytes, although all cell types and timepoints enrich for TNFA signalling via NF-κB (Figure 6).

Notably, irradiated SASPs significantly enriched for the epithelial-mesenchymal transition (EMT) pathway (SI Figure 19b). EMT is a critical process for cellular reprogramming [9, 13]. Importantly, senescent cells stimulate cellular reprogramming [14]. Considering that p16 and IL6 have specifically been linked to facilitating reprogramming [15], it is plausible that heterogeneous SASP secretions help modulate EMT transitions, facilitating reprogramming, although more research must be done.

### 3.2.4 Apoptosis

As both CS and CQ states involve the upregulation of anti-apoptotic pathways, understanding how pro- and anti-apoptotic SRMs are regulated may lead to better senolytics [16, 17]. While strategies have been developed to target these anti-apoptotic pathways in order to eliminate senescent cells, the heterogeneous nature of senescent phenotypes results in senolytics selectively targeting some populations of senescent cells over others [18-20].

We identified heterogenous induction of pro- and anti-apoptotic transcripts up- or down-regulated in fibroblasts induced into CQ and CS (Figure 4a). Some of these important apoptosis-associated genes – such as *BBC3* which encodes PUMA, alongside *PMAIP1* which encodes NOXA – also showcased heterogeneous temporal changes in the irradiation-induced senescent DEGs, dependent on cell type (Figure 6b). Furthermore, overexpressed fibroblast and melanocyte CS DEGs significantly enriched for the apoptosis pathway, whereas keratinocytes did not, further highlighting cell type-specific differences in how 'senescent' cells modulate apoptosis.

### 3.3.1 Transcription Factors

p53 is a key TF regulating various stress responses [21], and we found it was significant downregulated in a context-dependent manner (Figure 4a and 6b). This is not necessarily surprising given that p53 is known to degrade following activation of the unfolded protein response, which is itself activated in 'senescence' [22, 23]. Nonetheless, the heterogeneous expression of p53 and regulators of p53 like MDM2 can directly modulate the SASP and likely play a role in how other SRMs are regulated [24, 25].

Other TFs were significantly dysregulated depending on context, including *ATF4* – the main effector of the integrated stress response (ISR), which tailors how cells respond to stress (Figure 4a and 6b) [26]. Importantly, the ISR modulates various SRMs related to inflammation and autophagy [27-29].

Various pathways that regulate the SASP are known to converge on NF-κB and c/EBPβ [30-35]. Both TFs have been linked to regulating inflammation in general [36-38]. We found that *CEBPB* was overexpressed in SIPS and OIS arrest-DEGs, but not RS (Figure 4a). This may be one potential reason why RS appeared less proinflammatory than SIPS and OIS.

### 3.3.2 Chromatin Rewiring

One intriguing pattern is the universal downregulation of *LMNB1* across arrest-DEGs, alongside keratinocyte and fibroblast temporal DEGs. We found that melanocytes do not downregulate *LMNB1* until later timepoints (Figure 4a and 6b). This is of particular interest because global downregulation of *LMNB1* is a key event in 'senescence' induction [39-41], which may in part explain transcriptomic differences between the irradiation-induced melanocyte CS programme compared to other cell types (Figure 6a). Furthermore, *HMGA* transcripts were significantly overexpressed in OIS fibroblasts as expected, further suggesting chromatin rewiring and SAHF formation in this context (Figure 4a) [42].

### 3.3.4 Metabolism

- Nucleotide metabolism: Downregulation (partially significant) across all five arrest conditions, together with the folate cycle (synthesizing dTMP), is consistent with lack of DNA replication in arrested cells in general, and in particular with the promotion of CS through nucleotide synthesis inhibition [43-45].
- Energy metabolism, specifically glycolysis, TCA cycle and oxidative phosphorylation, overall show a trend of downregulation in CQ but sustained activity in CS. This might correspond to overall reduced energy demands in CQ as a result of the cell arrest, which in CS might be negated by the increased metabolic demand to maintain CS-specific phenotypes like the SASP [46].
- Inflammation and SASP: COX-2 – encoded by the *PTGS2* gene – plays a crucial role in prostaglandin metabolism [47]. Prostaglandins and their metabolic pathway have been shown to both promote the senescent phenotype and influence the composition of the

SASP [48, 49]; with COX-2 controlling the expression of SASP components, including IL6, IL8, CXCL1, and SAA1. *PTGS2* was significantly upregulated in OIS and SIPS, but not RS or CQ – arrest conditions that did not significantly overexpress various pro-inflammatory SASP proteins (Figure 4a). As can further be seen in the eicosanoid COX map (SI Figure 15), the COX-2 product prostaglandin H2 (PGH2) is further converted to prostaglandin E2 (PGE2) via *PTGES*. *PTGES* is significantly upregulated in SIPS, but significantly downregulated in RS, which may further explain why RS samples were less proinflammatory (Figure 4a).

- NAD salvage: *NAMPT* — a rate-limiting NAD enzyme —has been shown to inhibit CS in rat mesenchymal stem cells [50]. *NAMPT* was upregulated across most arrest conditions — except RS and early fibroblast irradiated samples — but was downregulated in melanocytes across all time points (SI Table 2, 28). This is in line with previous publications that show that *NAMPT* is downregulated in RS and modulates inflammation [51], although it is unclear why melanocytes downregulate this enzyme.

CS and CQ show metabolic alterations that can be related to specific arrest conditions and related non-metabolic processes (e. g. SASP activity and components). However, the statistical signals of individual pathways were much weaker than for many of the non-metabolic pathways studied in this work. This might be related to several challenges in the transcriptomic analysis of metabolic pathways, outlined below.

The small size of some metabolic pathways studied here poses inherent challenges in achieving statistical significance in ORA. Additionally, some reactions are catalysed by isozymes, of which only a subset might be expressed or perturbed between two conditions, potentially further weakening statistical signals. Moreover, in a bidirectional pathway like glycolysis/gluconeogenesis, many enzymes are shared between the two directions, such that up- or downregulation of these shared enzymes on their own does not provide information on the relevant directionality. Finally, relevant biological signals may not lie in whole-pathway-level up- or downregulations but in specific differential expression patterns within the pathway, such as directing metabolic flux toward particular pathway intermediates as touched upon in Section 3.3.4 for the eicosanoid COX map (SI Figure 15).

The purely transcriptomic scope of this study means that any downstream features on the proteomic or metabolomic level are not directly accessible. For instance, it was recently found that the glycolytic enzyme pyruvate kinase M2 (*PKM2*) forms aggregates in senescent cells from mice, decreasing glycolytic flux [52]. In another example, activity of the pentose phosphate pathway enzyme glucose-6-phosphate dehydrogenase (*G6PDH*) is inhibited by association with cytoplasmic p53 in unstressed cells, which is released under cellular stress conditions to thereby activate *G6PDH* [53]. Furthermore, perturbations at the enzyme and metabolite levels do not necessarily correlate, as shown by Wu et al. in upper glycolysis of a Dox-induced CS model [54].
