## Supplemental figures for "Mosaic Regulation of Stress Pathways Underlies Senescent Cell Heterogeneity"

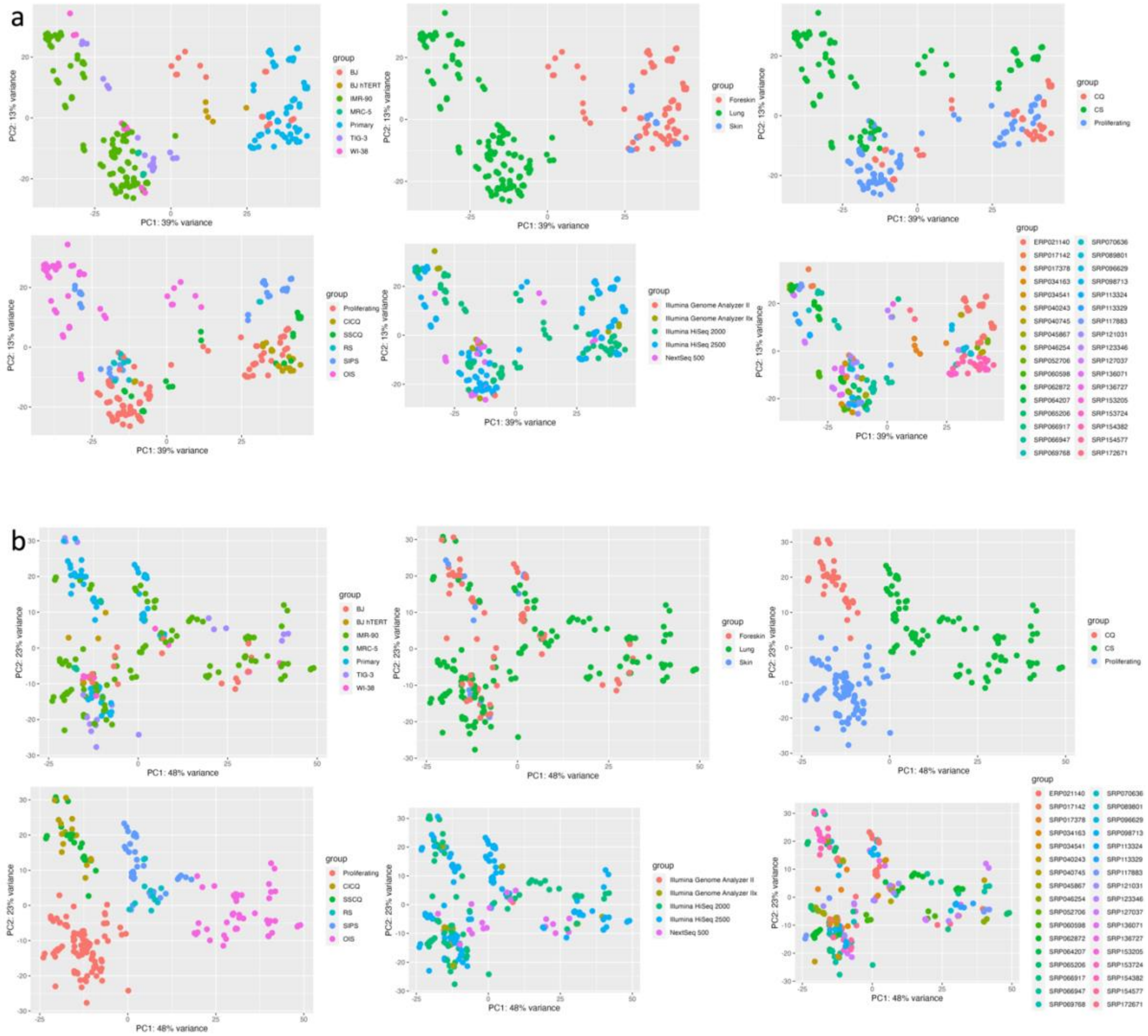


*SI Figure 1. PCA of arrest samples (a) before and (b) after removing the study batch effect. The following covariates are shown, from left to right: cell line, tissue, cell state, cell sub-state, sequencing platform, and SRA accession. CICQ: contact-inhibited CQ; SSCQ: serum-starved CQ.*

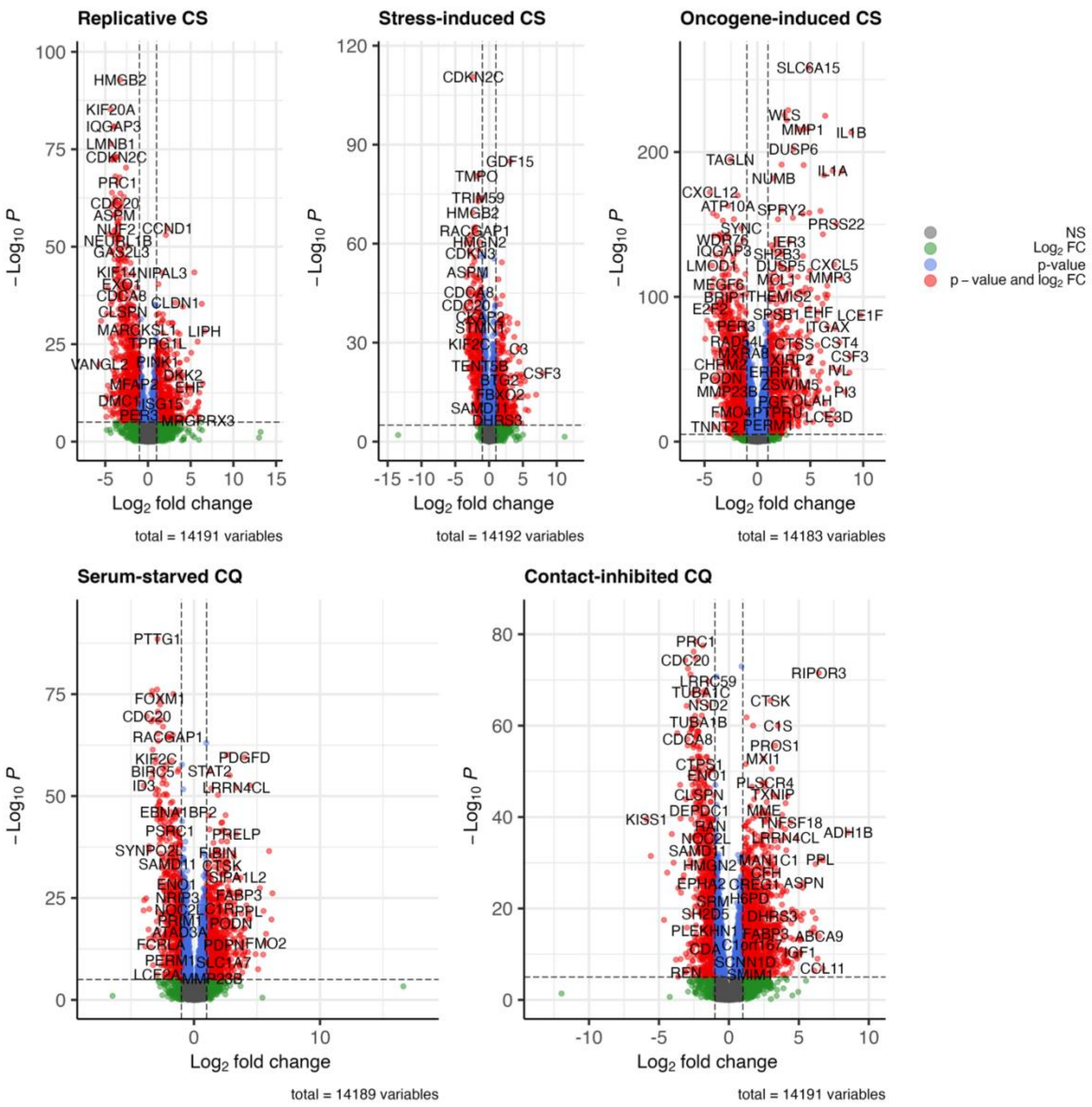


*SI Figure 2. Volcano plots of DEGs. Genes in red show significant p-value and $log_2$FC compared to proliferating controls ($p<0.05$ and $|log_2FC|>log_2(1.5)$). Genes in blue had significant p-value changes but not $log_2$FC changes, while genes in green had significant $log_2$FC changes but not significant p-value changes.*

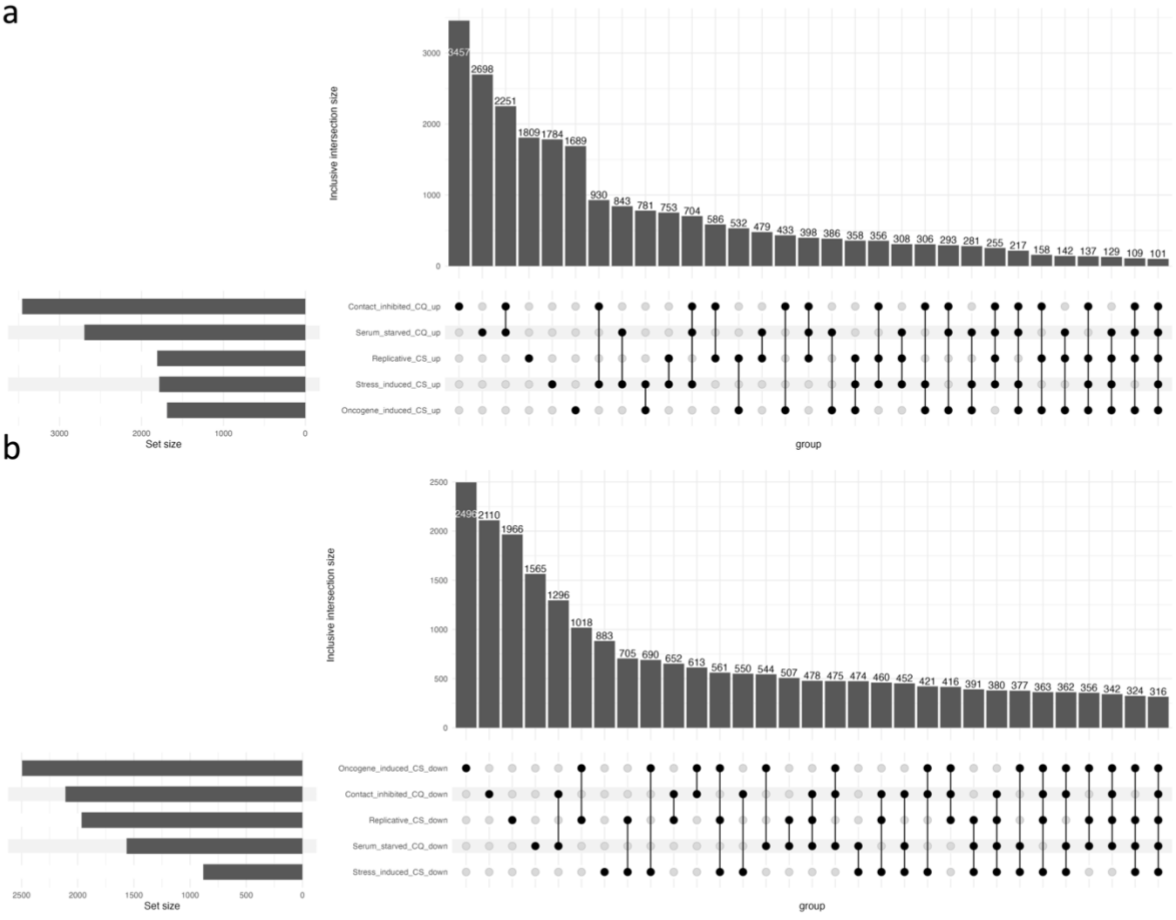


*SI Figure 3. Upset plots of DEGs a) upregulated and b) downregulated in the same direction compared to proliferating controls.*

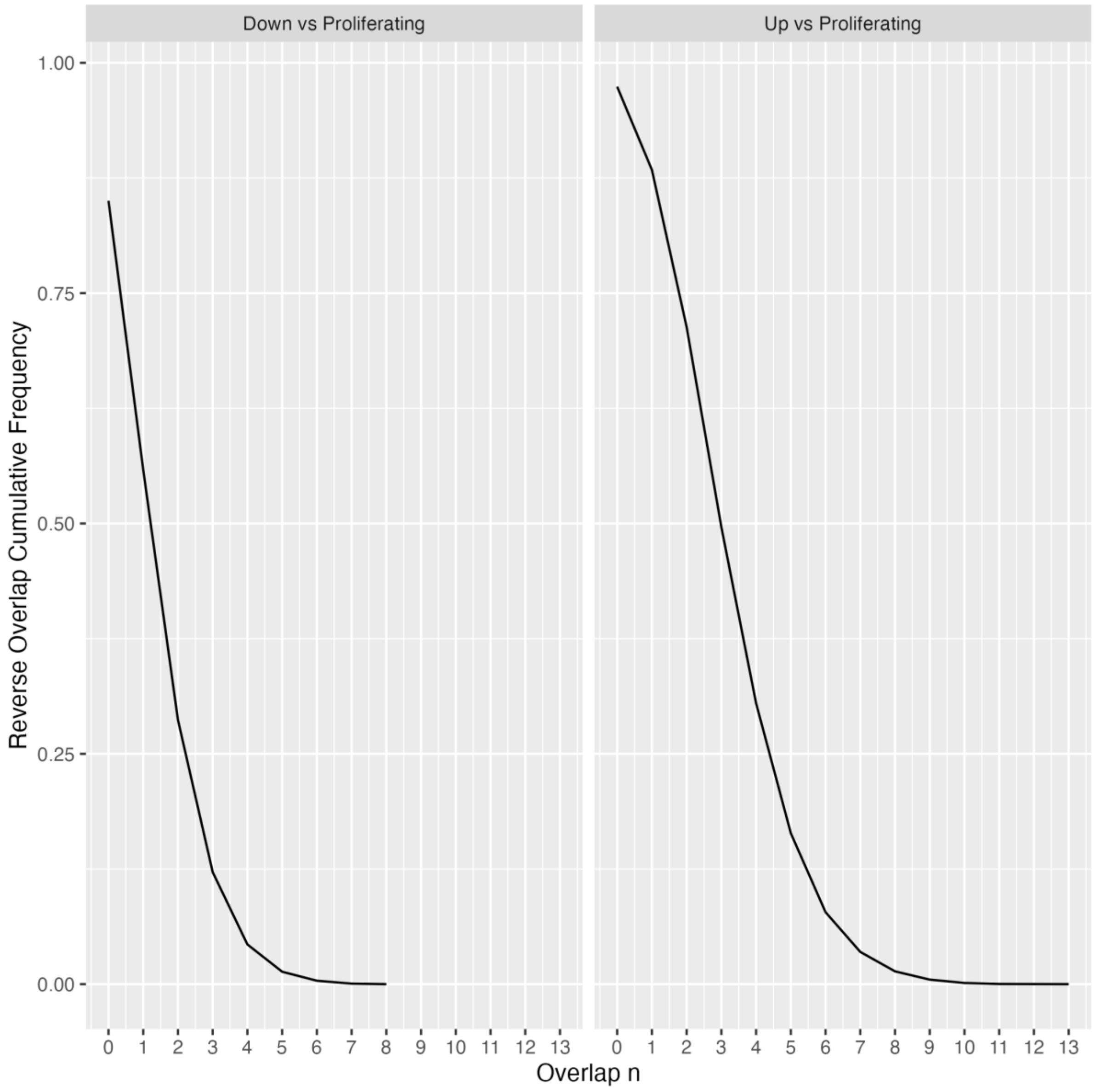


*SI Figure 4. Reverse cumulative frequency (y-axis) of number of simulated gene overlaps (x-axis) under- and overexpressed across all five arrest conditions in 10,000 simulations.*

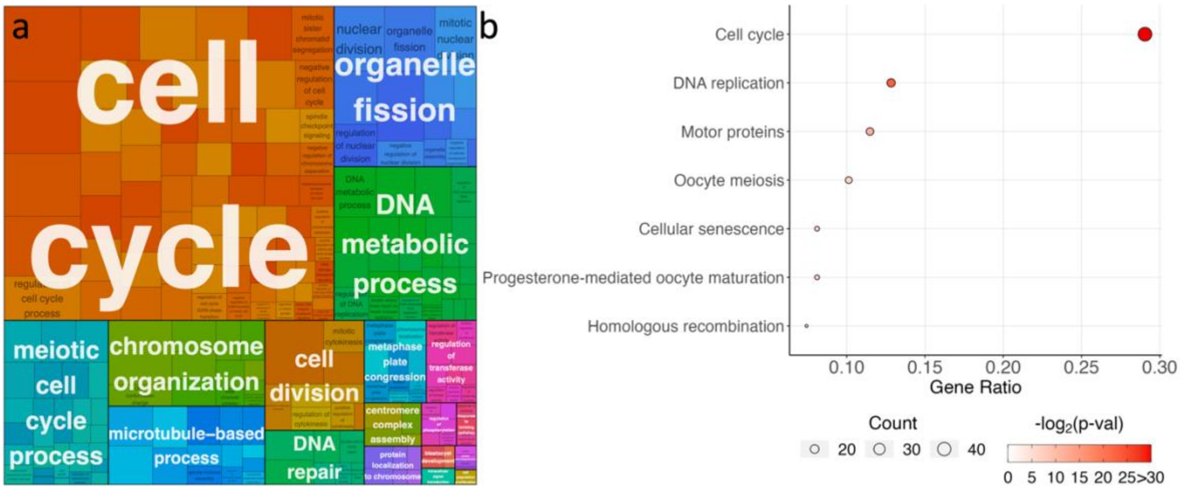


*SI Figure 5. Enriched a) GO and b) KEGG pathways amongst the 316 shared underexpressed DEGs, using all genes underexpressed across all five arrest conditions as an enrichment background.*

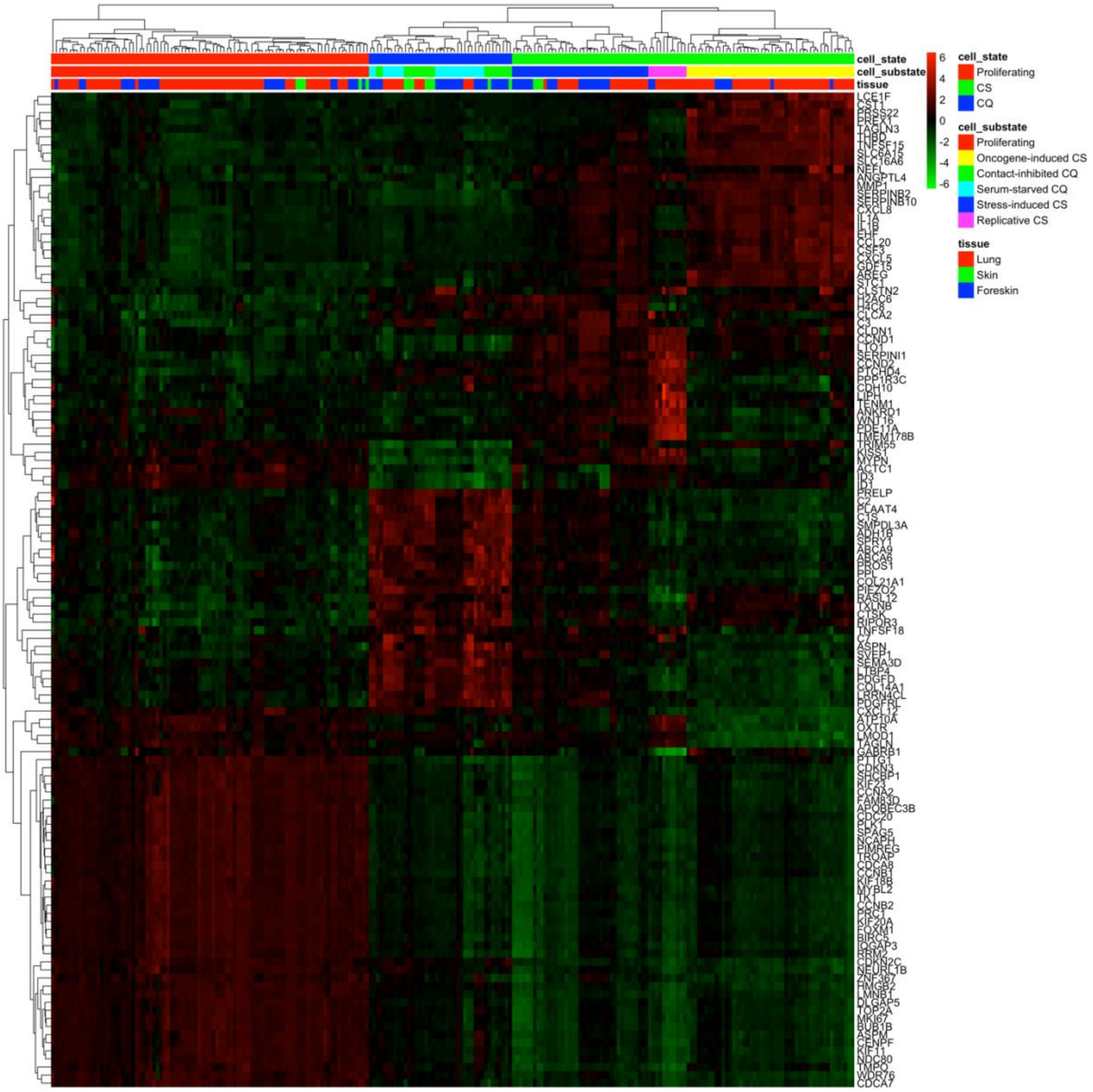


*SI Figure 6. Unsupervised hierarchal clustering of proliferating and arrested samples using the top 15 DEGs for each condition. Some of the top DEGs were shared between arrest conditions.*

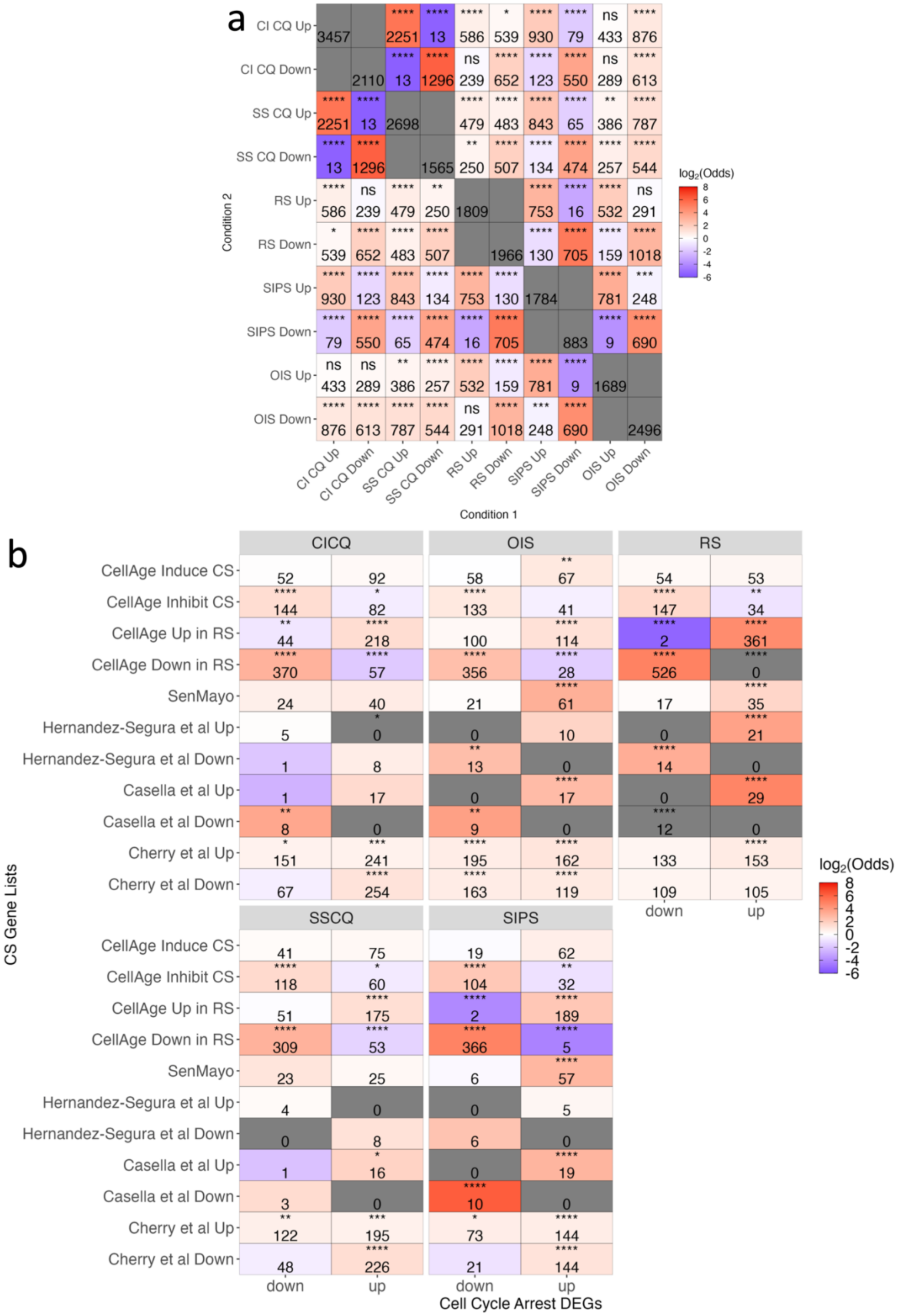


*SI Figure 7. a) Overlap between overexpressed and underexpressed RS, SIPS, OIS, contact-inhibited (CI) and serum-starved (SS) CQ DEGs between each arrest condition. Note that overlaps are*

*duplicated for easier visualisation. Grey tiles indicate self-overlaps, and numbers in grey boxes indicate the total number of DEGs for the given condition. b) Overlap between CS- and CQ-DEGs and signatures of CS, including drivers of CS and signatures of RS from CellAge and CS signatures derived from various studies. Red tiles indicate more overlaps than expected by chance (positive $log_2$(odds ratio)), whereas blue tiles indicate less overlaps than expected by chance (negative $log_2$(odds ratio)). Grey tiles represent no overlaps between conditions (infinite odds ratio). Significance was assessed using a two-tailed Fisher's exact test with Bonferroni correction (ns – not significant, * - p<0.05, ** - p<0.01, *** - p<0.001, **** - p<0.0001).*

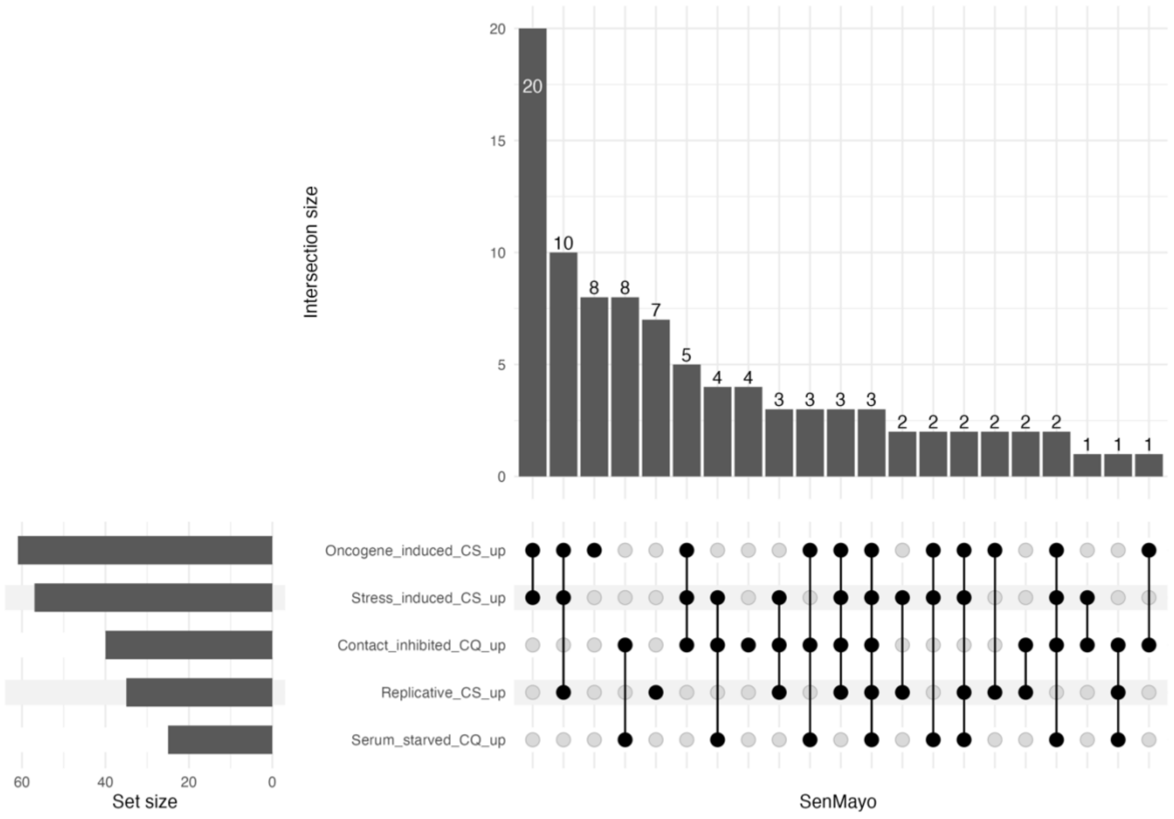


*SI Figure 8. Upset plot of cell cycle arrest DEGs upregulated in the SenMayo dataset.*

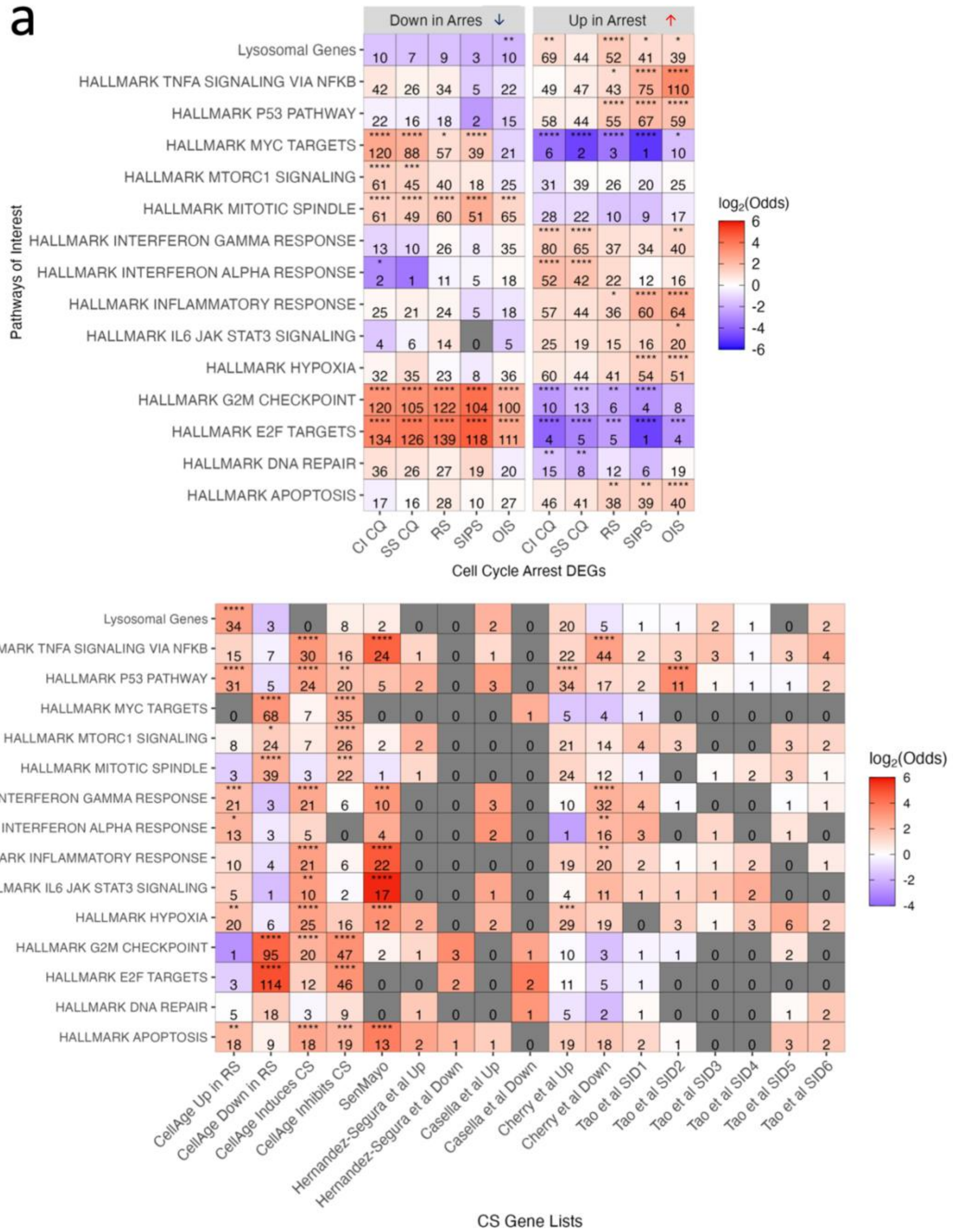


*SI Figure 9. a) Overlap between overexpressed and underexpressed RS, SIPS, OIS, CI and SS CQ DEGs and lysosomal genes and MSigDB stress-associated pathways. b) Overlap between CS-associated genelists and the aforementioned pathways. Note that while overlaps were performed for the entire MSigDB, only some pathways are shown due to space constraints.*

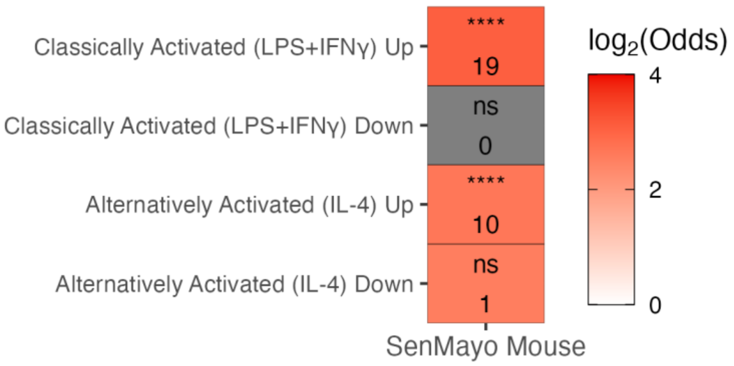


*SI Figure 10. Overlap between macrophage DEGs and SenMayo mouse, using all protein-coding mouse genes as an overlap background.*

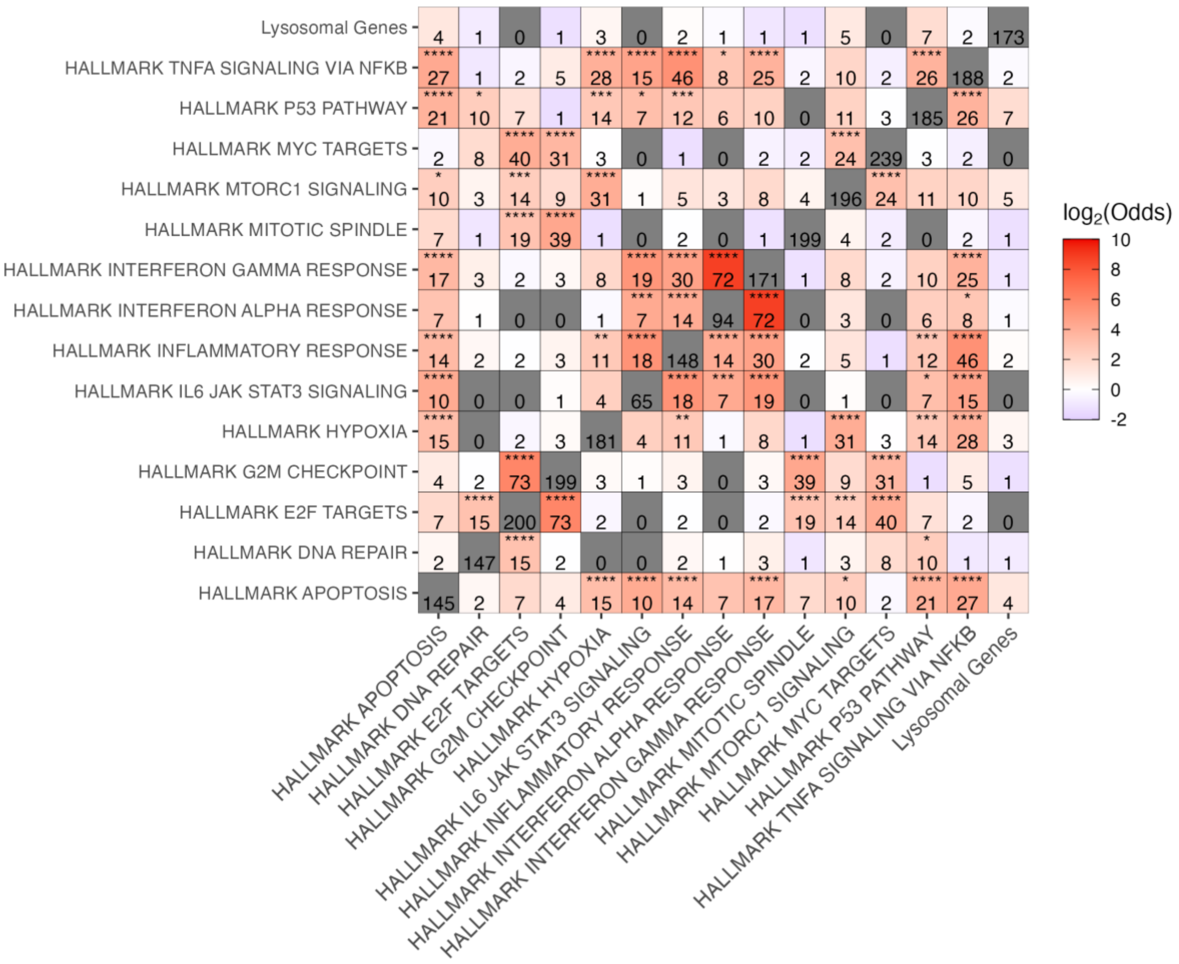


*SI Figure 11. Overlap between MSigDB stress-associated pathways. Note that overlaps are duplicated for easier visualisation. Grey tiles indicate either no overlaps, or self-overlaps, and numbers in tiles indicate the total number of genes for the given overlap or pathway. Red tiles indicate more overlaps than expected by chance (positive $log_2$(odds ratio)), whereas blue tiles indicate less overlaps than*

*expected by chance (negative log$_2$(odds ratio)). Note that while overlaps were performed for the entire MSigDB, only some pathways are shown due to space constraints.*

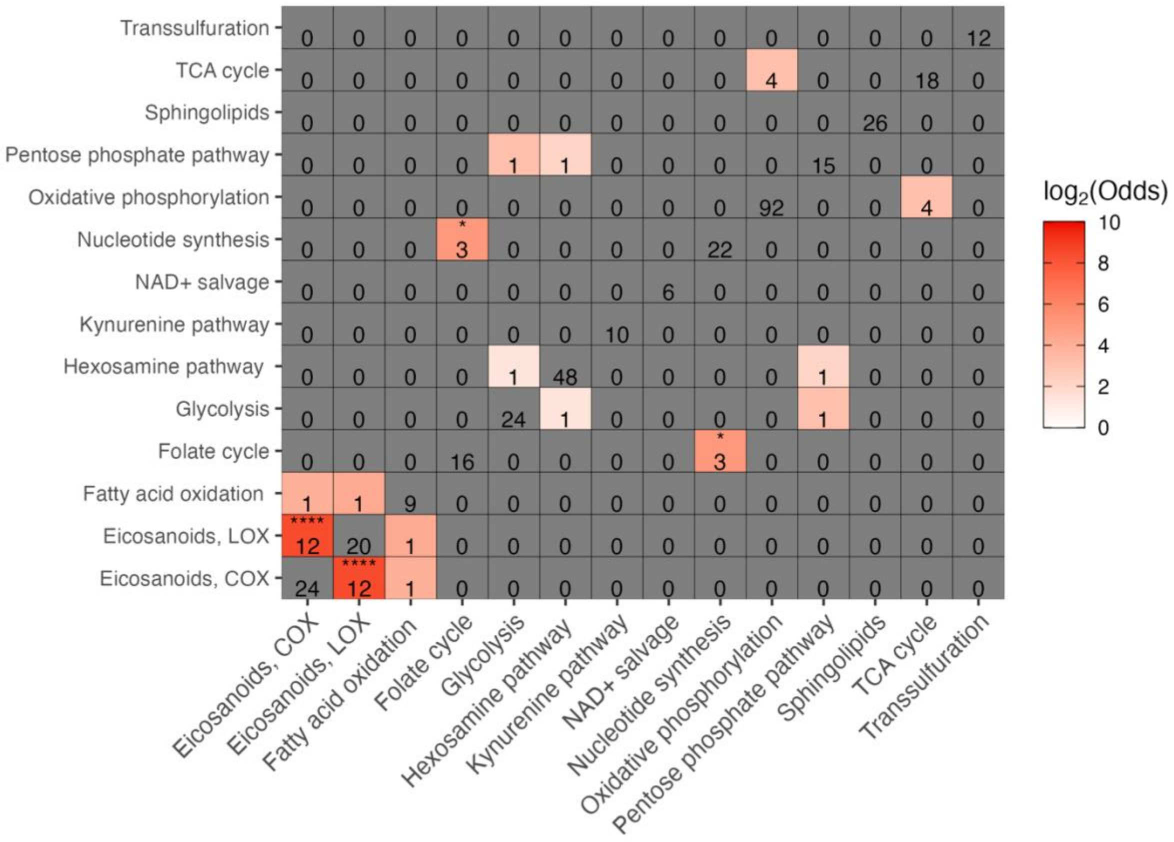


*SI Figure 12. Shared genes between metabolic pathways.*

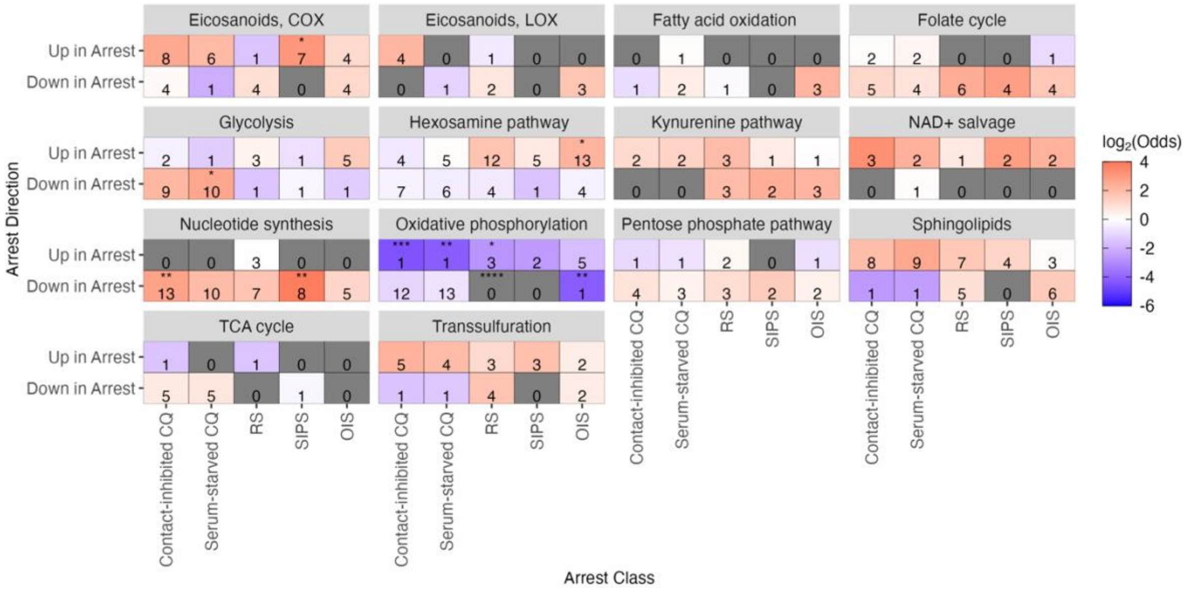


*SI Figure 13. Overlap between over- and underexpressed arrest-DEGs and metabolic enzymes from various pathways. Significance assessed using a negative binomial distribution with BH correction and $|log_2(FC)|>log_2(1.5)$). Maximum $log_2FC$ was capped at 6 to visualise differences more clearly between conditions. * - p<0.05, ** - p<0.01, *** - p<0.001, **** - p<0.0001.*

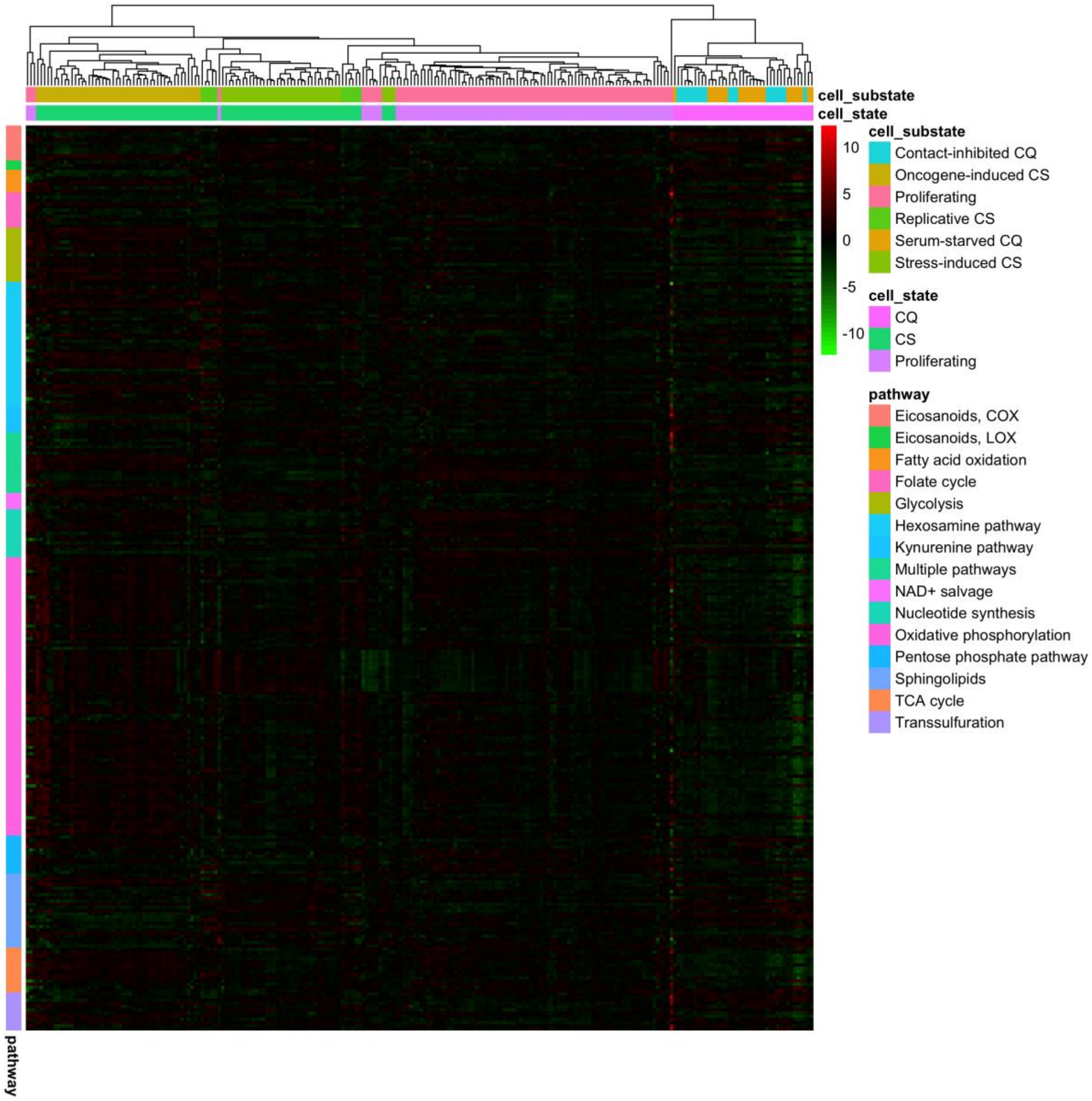


*SI Figure 14. Hierarchal clustering of arrested and proliferating fibroblasts based on gene expression from metabolic pathways. Genes are grouped by pathway, while samples were clustered based on pairwise distances between samples using scaled normalised gene expression profiles.*

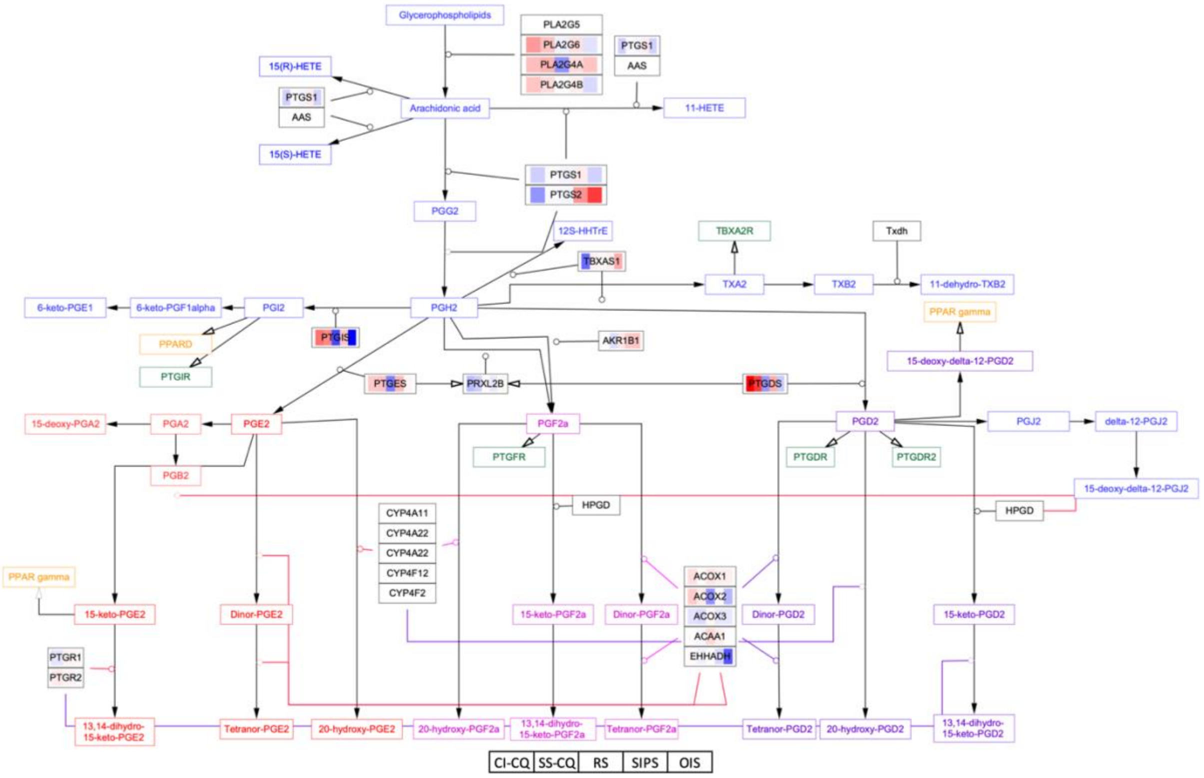


*SI Figure 15. Log$_2$(FC) of relevant arrest DEGs mapped to the Eicosanoid metabolism via cyclooxygenases WikiPathway with ID number WP4347. Tiles in red indicate overexpression, whereas blue tiles indicate underexpression compared to proliferating controls. Each gene shows 5 conditions: contact-inhibited CQ, serum-starved CQ, RS, SIPS, and OIS.*

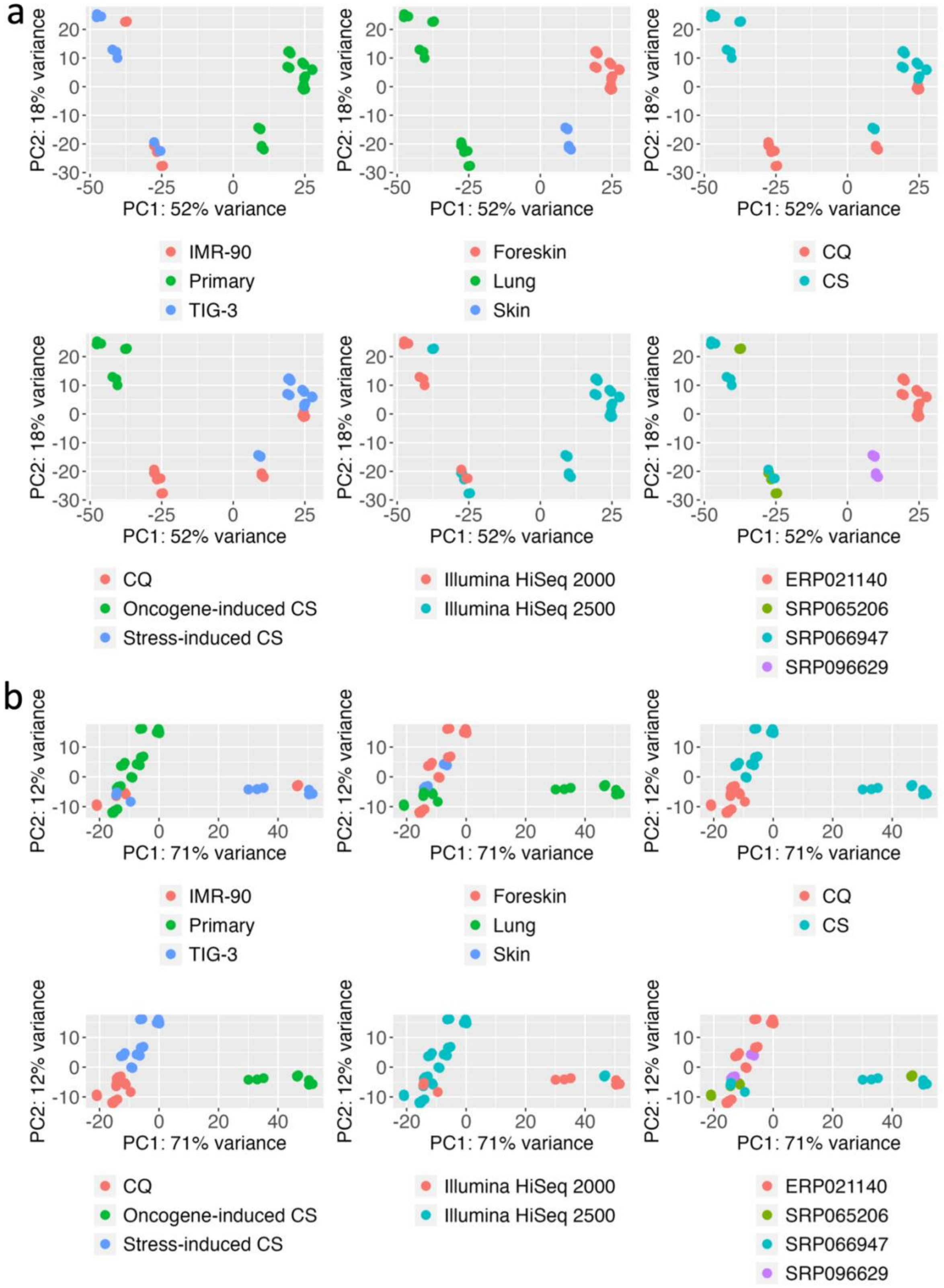


*SI Figure 16. PCA of SIPS, OIS, and CQ samples (a) before and (b) after removing the study batch effect. The following covariates are shown, from left to right: cell line, tissue, cell state, cell sub-state, sequencing platform, and SRA accession.*

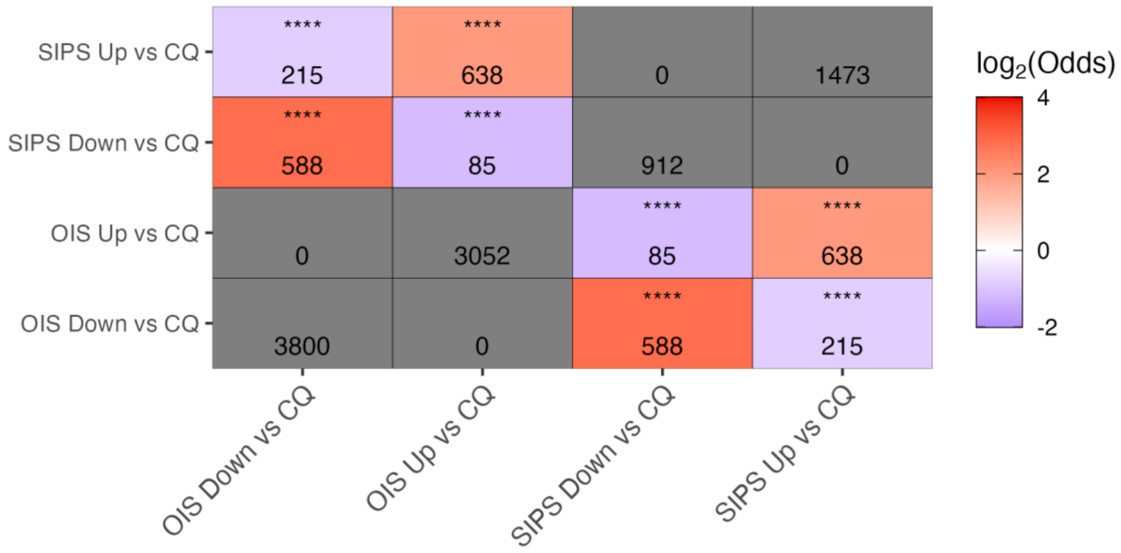


*SI Figure 17. Overlap between OIS and SIPS DEGs, by direction, generated against quiescent cells.*

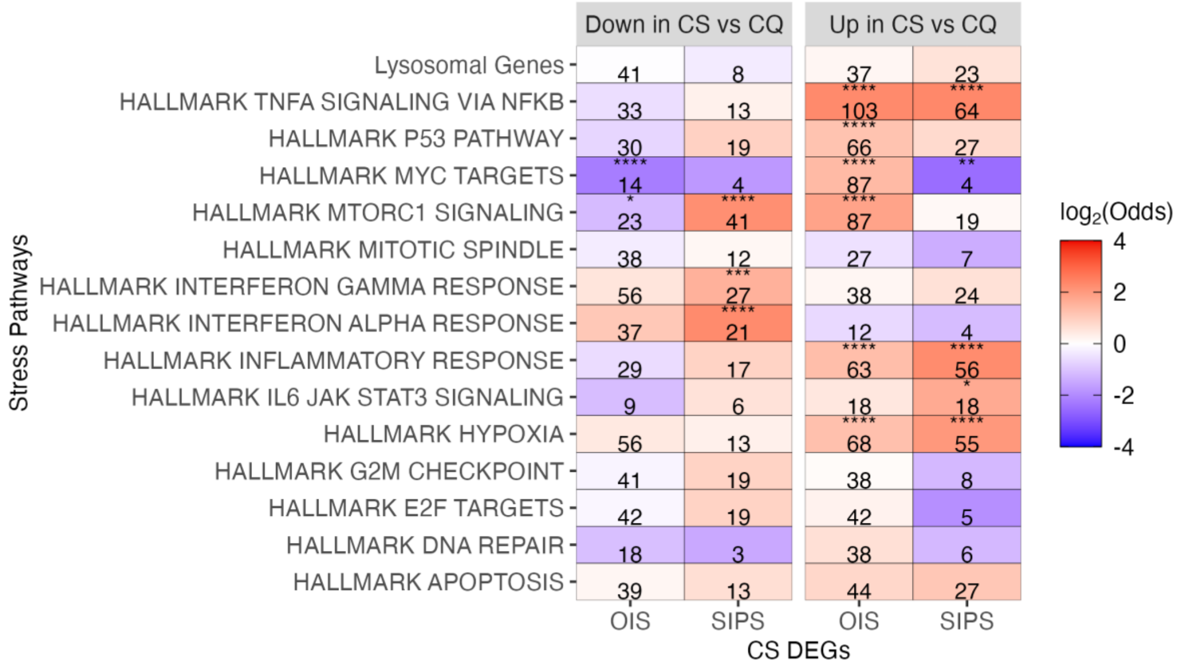


*SI Figure 18. Overlap of OIS and SIPS DEGs, by direction, generated against quiescent cells, with various stress response pathways. Note that while overlaps were performed for the entire MSigDB, only some pathways are shown due to space constraints.*

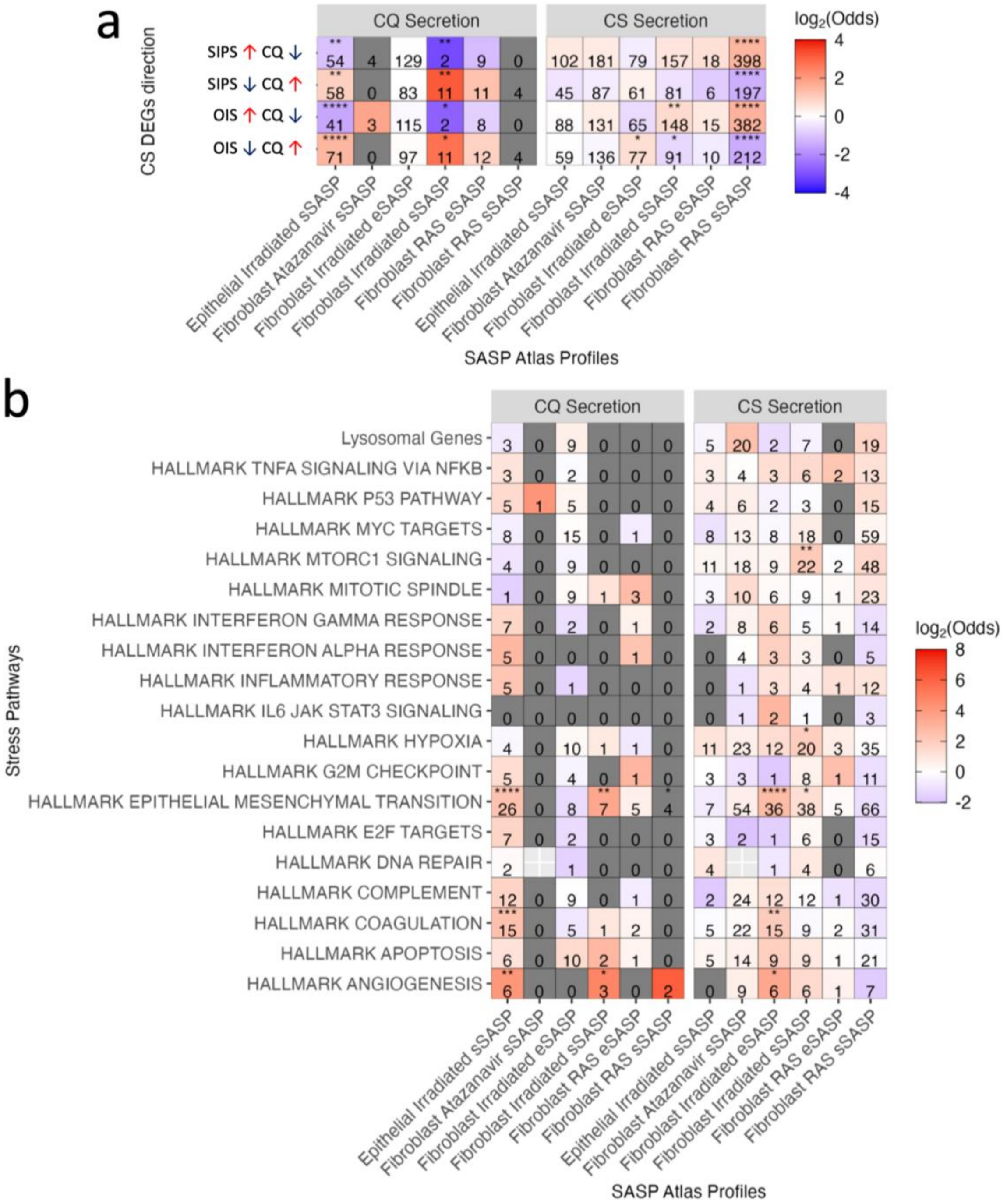


*SI Figure 19. a) Overlap between OIS and SIPS DEGs generated against CQ samples, and SASP atlas proteins, using the intersect of genes expressed in the recount3 data and proteins in the SASP atlas as the background. 'CQ Secretion' refers to proteins that were secreted significantly more in serum-starved quiescent samples compared to the given senescence conditions. b) Overlap between SASP atlas profiles and various pathways of interest. Note that while overlaps were performed for the entire MSigDB, only some pathways are shown due to space constraints.*

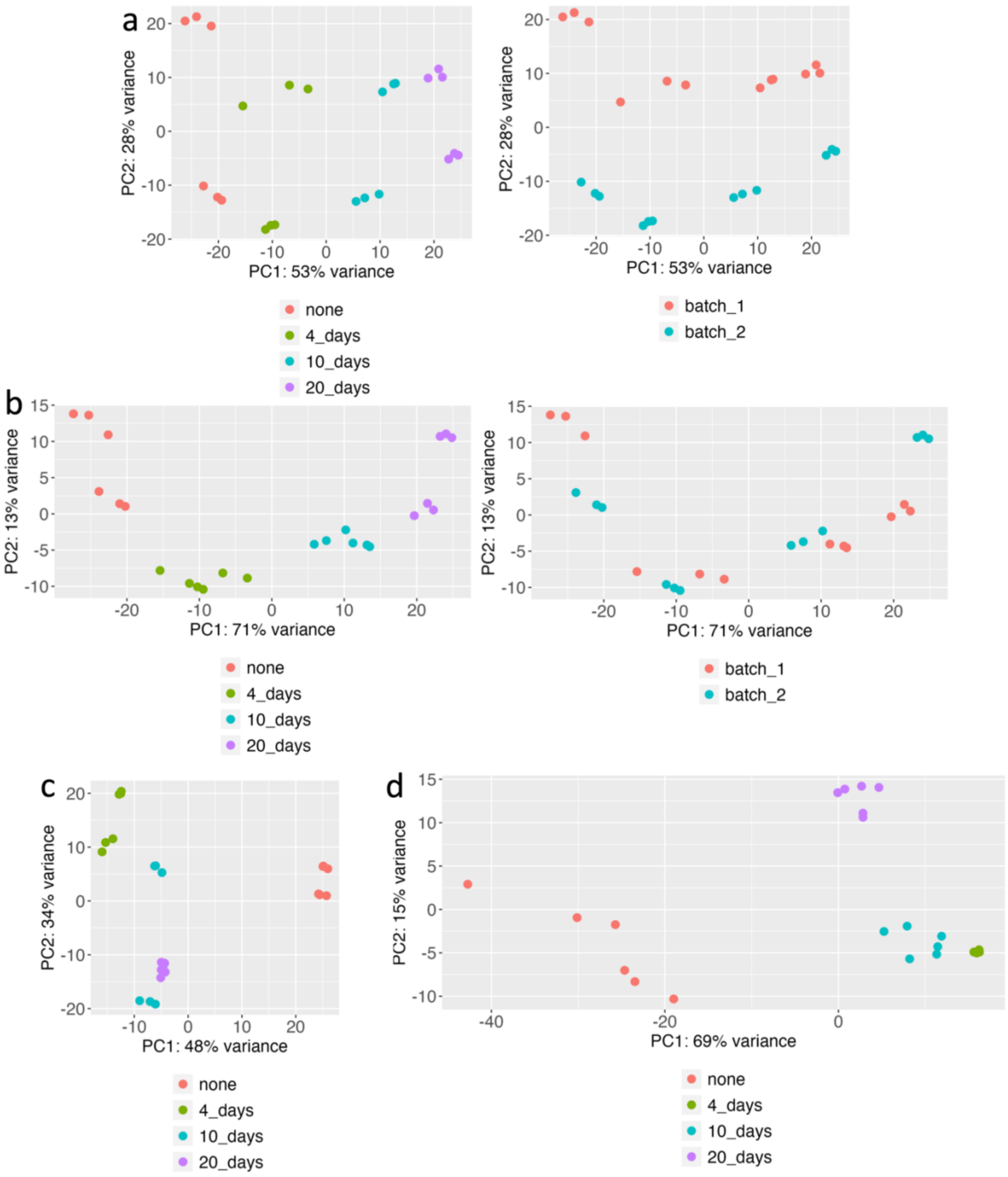


*SI Figure 20. PCA plots of temporal samples for keratinocytes (a) before and (b) after batch correction, (c) fibroblasts, and (d) melanocytes. 'None' refers to proliferating controls. The top 500 most variable genes were used for clustering.*

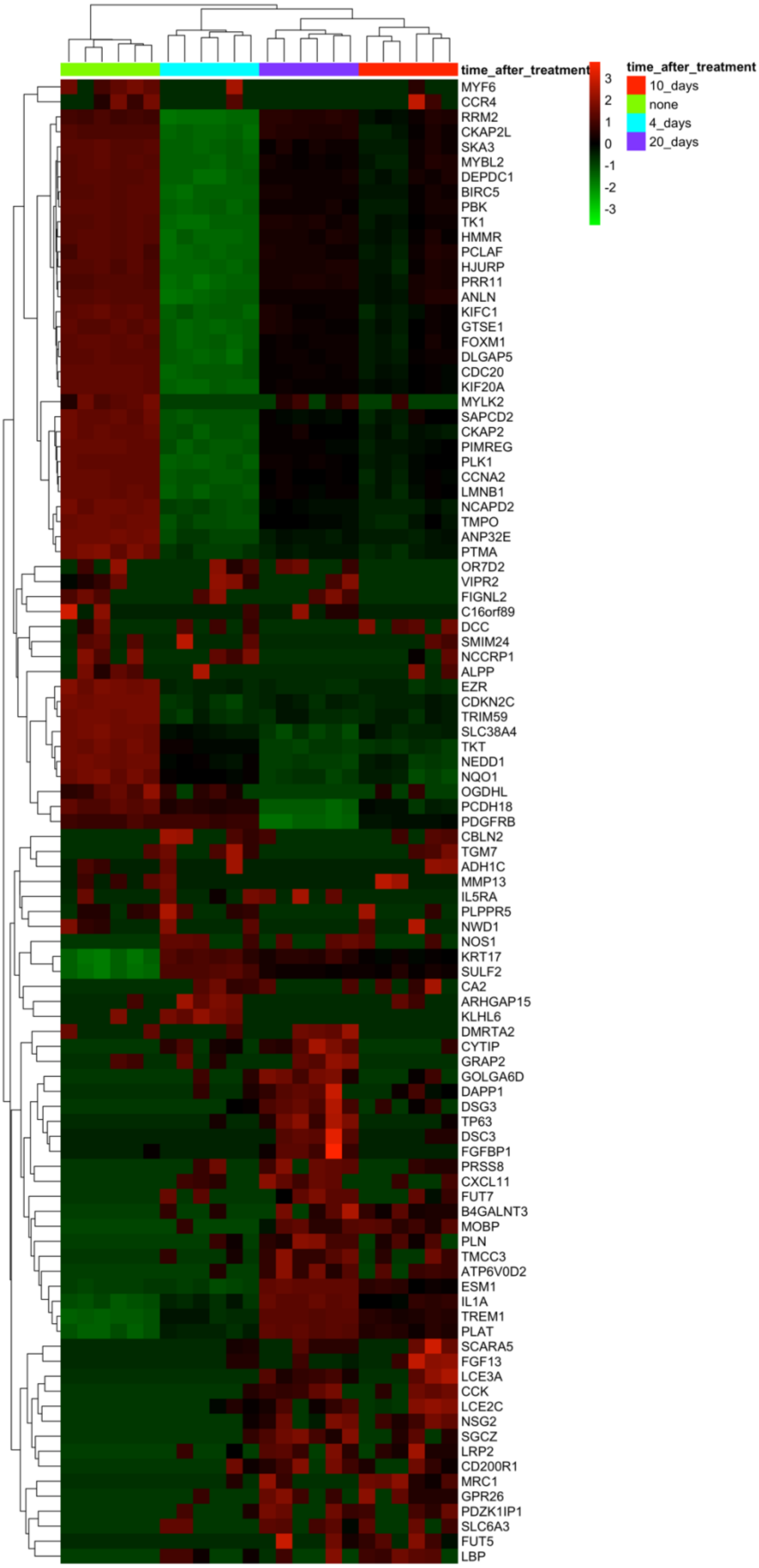


*SI Figure 21. Unsupervised hierarchal clustering of proliferating and arrested fibroblast samples using the top 25 DEGs from each condition.*

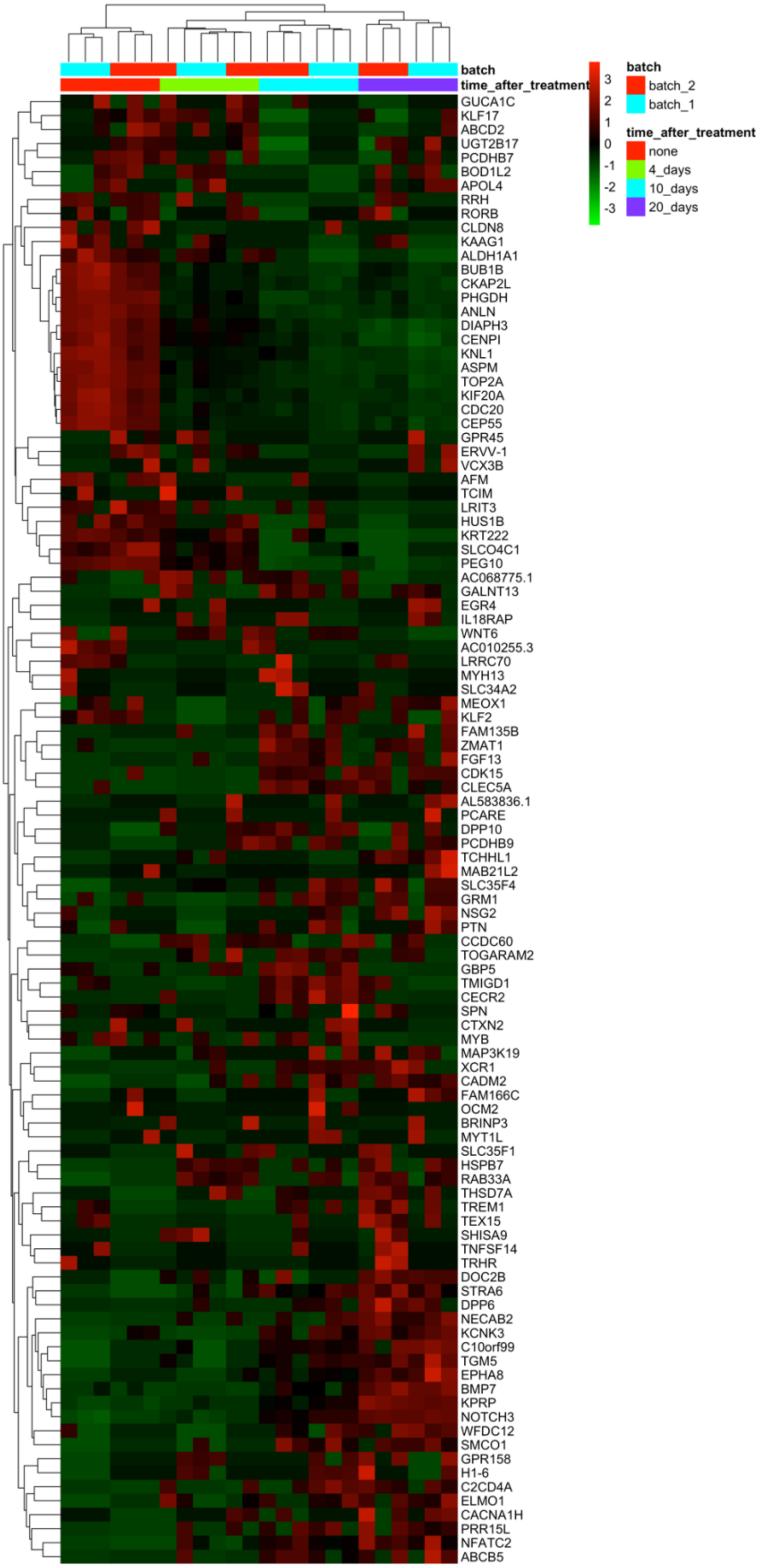

*SI Figure 22. Unsupervised hierarchal clustering of proliferating and arrested keratinocyte samples using the top 25 DEGs from each condition.*

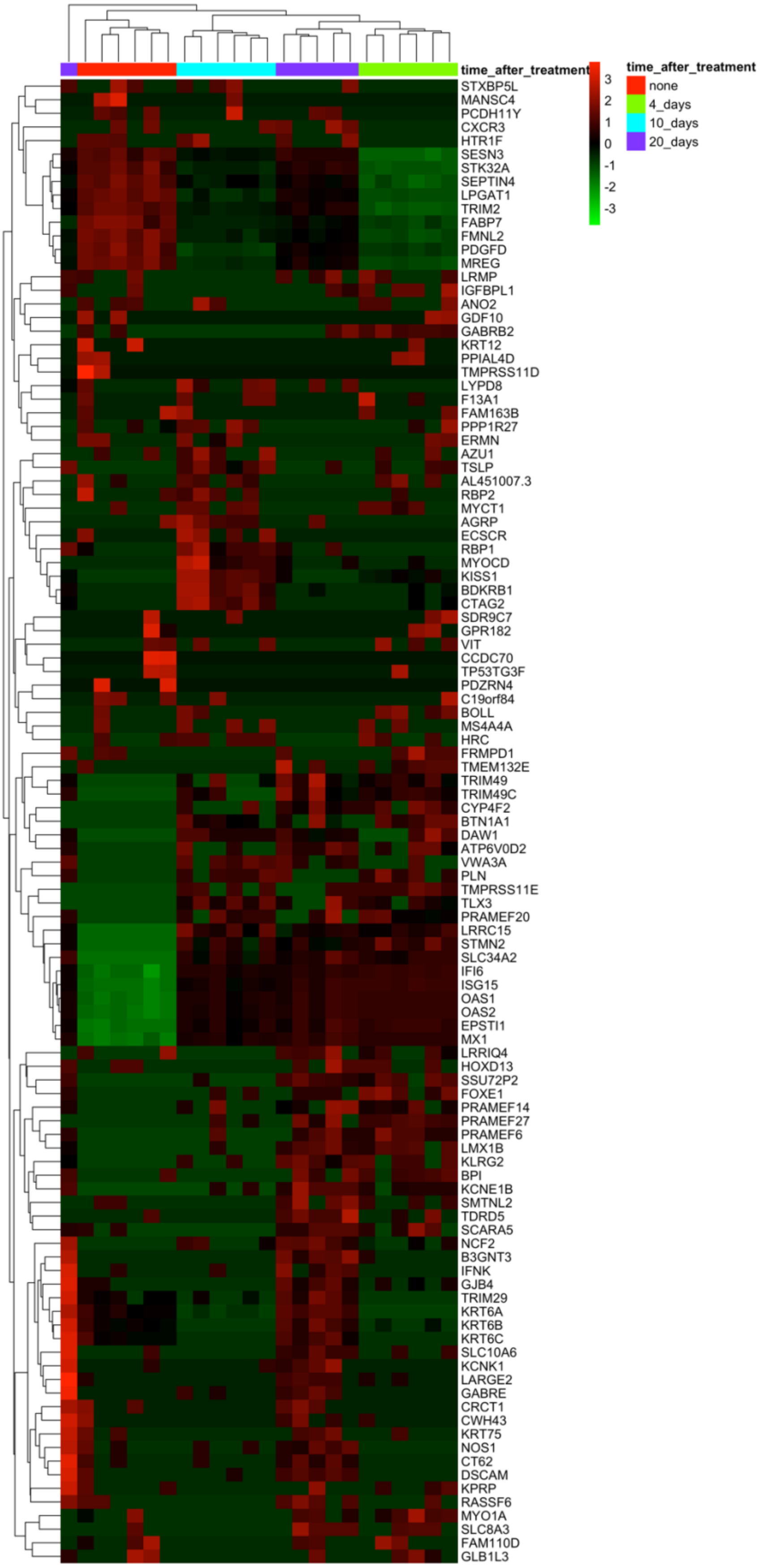


*SI Figure 23. Unsupervised hierarchal clustering of proliferating and arrested melanocyte samples using the top 25 DEGs from each condition.*

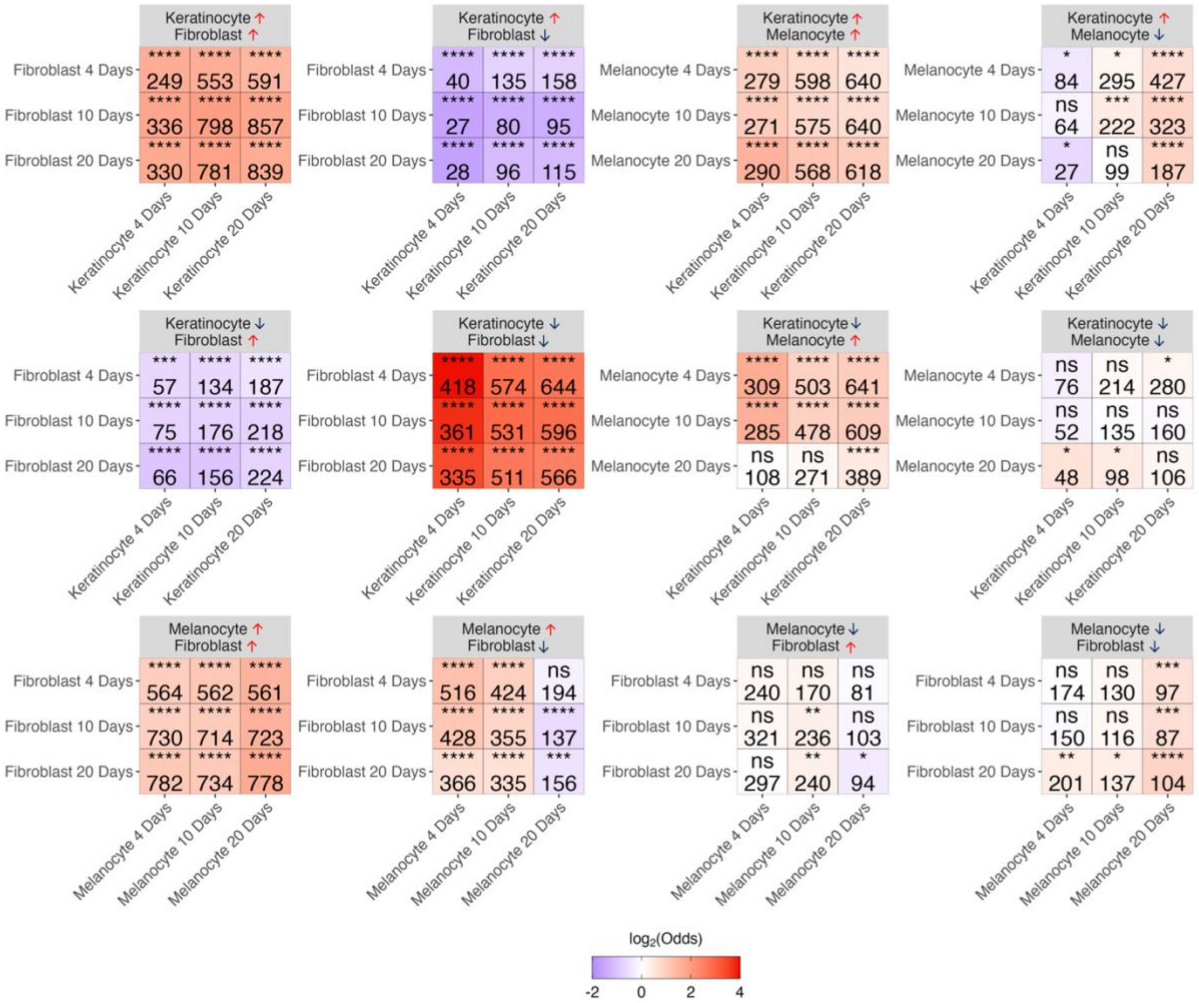


*SI Figure 24. Overlap of DEGs between cell types by days post-irradiation. Genes expressed in either cell type were used as the background.*

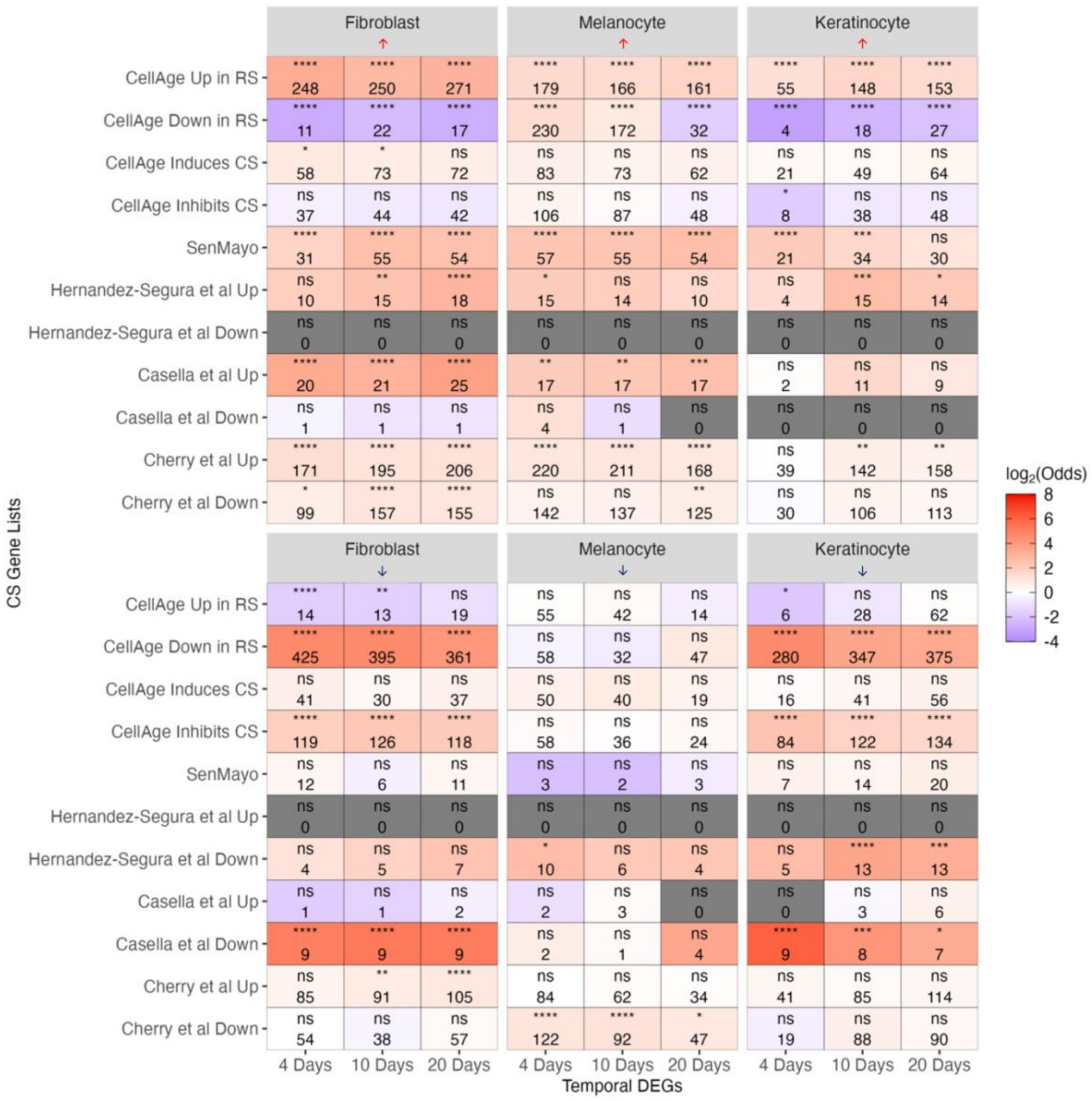


*SI Figure 25. Overlap between DEGs by day and cell type between various CS databases.*

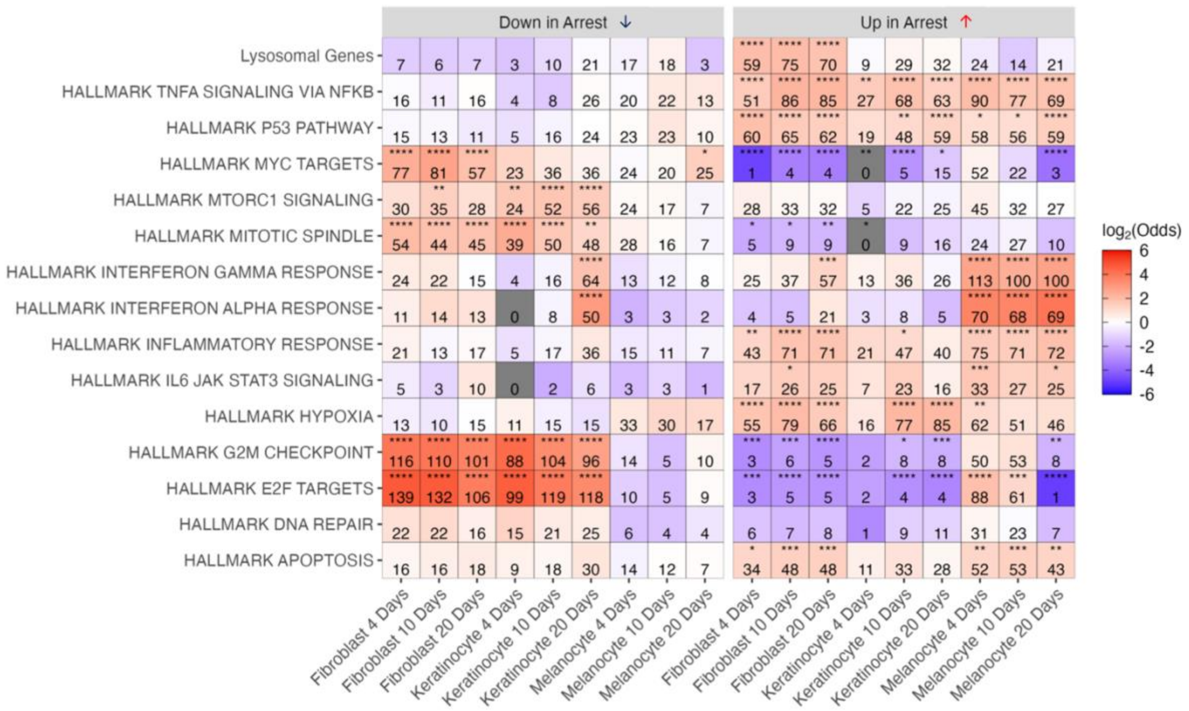


*SI Figure 26. Overlap between DEGs by day and cell type with various stress-associated pathways.*
